# Effects of periodic bottlenecks on the dynamics of adaptive evolution in microbial populations

**DOI:** 10.1101/2021.12.29.474457

**Authors:** Minako Izutsu, Devin M. Lake, Zachary W. D. Matson, Jack P. Dodson, Richard E. Lenski

## Abstract

Population bottlenecks are common in nature, and they can impact the rate of adaptation in evolving populations. On the one hand, each bottleneck reduces the genetic variation that fuels adaptation. On the other hand, each founder that survives a bottleneck can undergo more generations and leave more descendants in a resource-limited environment, which allows surviving beneficial mutations to spread more quickly. A theoretical model predicted that the rate of fitness gains should be maximized using ∼8-fold dilutions. Here we investigate the impact of repeated bottlenecks on the dynamics of adaptation using numerical simulations and experimental populations of *Escherichia coli*. Our simulations confirm the model’s prediction when populations evolve in a regime where beneficial mutations are rare and waiting times between successful mutations are long. However, more extreme dilutions maximize fitness gains in simulations when beneficial mutations are common and clonal interference prevents most of them from fixing. To examine the simulations’ predictions, we propagated 48 *E. coli* populations with 2-, 8-, 100-, and 1000-fold dilutions for 150 days. Adaptation began earlier and fitness gains were greater with 100- and 1000-fold dilutions than with 8-fold dilutions, consistent with the simulations when beneficial mutations are common. However, the selection pressures in the 2-fold treatment were qualitatively different from the other treatments, violating a critical assumption of the model and simulations. Thus, varying the dilution factor during periodic bottlenecks can have multiple effects on the dynamics of adaptation caused by differential losses of diversity, different numbers of generations, and altered selection.

**Significance:** Many microorganisms experience population bottlenecks during transmission between hosts or when propagated in the laboratory. These bottlenecks reduce genetic diversity, potentially impeding natural selection. However, bottlenecks can also increase the number of generations over which selection acts, potentially accelerating adaptation. We explored this tension by performing simulations that reflect these opposing factors, and by evolving bacterial populations under several dilution treatments. The simulations show that the dilution factor that maximizes the rate of adaptation depends critically on the rate of beneficial mutations. On balance, the simulations agree well with our experimental results, which imply a high rate of beneficial mutation that generates intense competition between mutant lineages.

## Introduction

Population sizes often fluctuate in time, sometimes greatly, with important effects on evolution (1, 2). A bottleneck occurs when a population’s size drops suddenly, typically leading to the loss of some genetic variants and a concomitant increase in frequency or even fixation of other variants. Bottlenecks occur in a variety of contexts, with some important distinctions including: (i) whether the survivors of a bottleneck are a random subset of the population or possess some particular trait (e.g., resistance to a selective agent); and (ii) whether a bottleneck and subsequent recovery are singular events or occur periodically (e.g., seasonally). Our focus here is on periodic bottlenecks in which the survivors are a random subset of the population. This situation arises with the serial transfer protocols used in many evolution experiments (3–5), and it is similar to the transmission of microbial pathogens and commensals between hosts (6–10). Therefore, the effects of periodic bottlenecks are important both for experimental evolution and for understanding the evolution of host-associated microbes.

In the case of extreme bottlenecks, a population’s mean fitness can decline over time, even in a constant environment. For example, in mutation-accumulation experiments, lineages are repeatedly propagated via single-cell bottlenecks (or full-sibling crosses for sexual organisms), thereby purging genetic variation and preventing natural selection from acting (except against lethal mutations, which are not observed and thus cannot be propagated). New mutations still arise in these experiments but, in the absence of selection, populations accumulate mutations without regard to their fitness effects; and because more mutations are deleterious than beneficial, the accumulation of these random mutations causes fitness to decline over time (11–13).

In this study, we are concerned with less severe bottlenecks that, nonetheless, can have important consequences for the dynamics of adaptation. In particular, we consider populations of asexual organisms that are propagated via periodic (e.g., daily) serial transfers into fresh medium. We consider cases in which the organisms can grow fast enough that, regardless of the dilution imposed, the population depletes the limiting resource before the next dilution takes place. The population thus reaches a final stationary-phase density prior to the next transfer, and we assume for simplicity that no death occurs. The situation described above closely matches the long-term evolution experiment (LTEE) with *Escherichia coli* that has been running for over 35 years and 75,000 generations (3, 13–17). Each day in the LTEE, the bacteria are diluted 100-fold into 10 mL of fresh medium, with glucose as the limiting resource. The glucose concentration allows the bacteria to reach a final density of ∼5 × 10^7^ cells per mL, or ∼5 × 10^8^ cells in total. After each dilution, the population then grows 100-fold to reach stationary phase. That regrowth corresponds to log_2_ 100 ≈ 6.6 doublings per day. In this medium, the ancestral bacteria grow with a doubling time of ∼1 h, and so they readily reach the final density well before the next dilution and transfer. Indeed, they could be diluted a million-fold and still achieve the 20 doublings required to reach stationary phase. Depending on the particular strain and culture medium, some bacteria might experience death after they exhaust their resources. However, in the case of the ancestral strain and culture conditions used in the LTEE, there is little or no cell death between transfers (18).

We ask the following question: What daily dilution factor, *D*, would maximize the rate of fitness gain? Changing the dilution factor has two opposing effects. On the one hand, more severe bottlenecks cause the loss of a higher proportion of beneficial mutations soon after they arise and while they are still at low frequencies, resulting in fewer surviving beneficial mutations. On the other hand, more severe bottlenecks increase the number of generations between successive transfers, which amplifies the growth advantage of those beneficial mutations that survive a bottleneck. For example, populations subjected to *D* = 2 have only 1 generation (i.e., one doubling) between transfers, while populations experiencing *D* = 8 have 3 generations per transfer and those propagated at *D* = 1000 have ∼10 generations between transfers. (More severe bottlenecks also mean that more cells are produced during the regrowth, with the potential for more new beneficial mutations at higher dilution factors. However, this effect cannot be greater than 2-fold, because at least as many new cells are produced in the last population doubling as in all previous doublings.) We consider the case where the fate of individuals during dilution is random; thus, the fitness effect of a mutation does not influence whether it survives the bottleneck. However, fitness differences matter during the regrowth between bottlenecks. An individual that has a beneficial mutation is expected to have more descendants than its parent before the next bottleneck, which increases the mutation’s probability of survival. Moreover, greater dilutions lead to more generations between transfers (assuming the dilutions are not so severe that they cause extinction), which allows the growth advantage of a beneficial mutation to compound faster on a per-transfer basis.

Wahl et al. (4) analyzed a mathematical model corresponding to this same scenario, and they derived the value of *D* that maximizes fitness gains given certain assumptions. According to their analysis, the supply of surviving beneficial mutations is maximized when *D* = *e*^2^ ≈ 7.4 (Fig. 1). Their analysis assumes that the evolving populations are in a strong-selection, weak-mutation (SSWM) regime, such that the fitness effects of beneficial mutations are large and beneficial mutations are rare (19–21). In this regime, the waiting time between the appearance of beneficial mutations bound for fixation dominates the dynamics, whereas the transit time for successful beneficial mutations is comparatively short (i.e., the time taken for a mutation to go from occurring in a single individual to being present in every individual). As a consequence, there is an absence of competition between multiple lineages that independently acquired beneficial mutations. Many theories of adaptation assume a SSWM regime. However, many microbial evolution experiments, including the LTEE, clearly violate the weak-mutation assumption. When beneficial mutations are common, the waiting times between beneficial mutations that survive genetic drift are short relative to the transit times required for their spread and eventual fixation. In asexual populations, this situation leads to clonal interference, in which multiple lineages with independent beneficial mutations must compete with one another and contend for fixation (22–28). Clonal interference causes those beneficial mutations that eventually fix to have larger effect sizes than those that would fix without that competition (29). Thus, the dilution factor that maximizes the rate of fitness increase when beneficial mutations are common remains unclear. Campos and Wahl (30) examined this case theoretically and in simulations that included both clonal interference and bottlenecks; that work assumed that each contending lineage had only a single beneficial mutation before one eventually fixed in the population. However, contending lineages can acquire multiple beneficial mutations before any of them become fixed if populations are large, have high beneficial mutation rates, or both (21, 31).

**Fig. 1.**
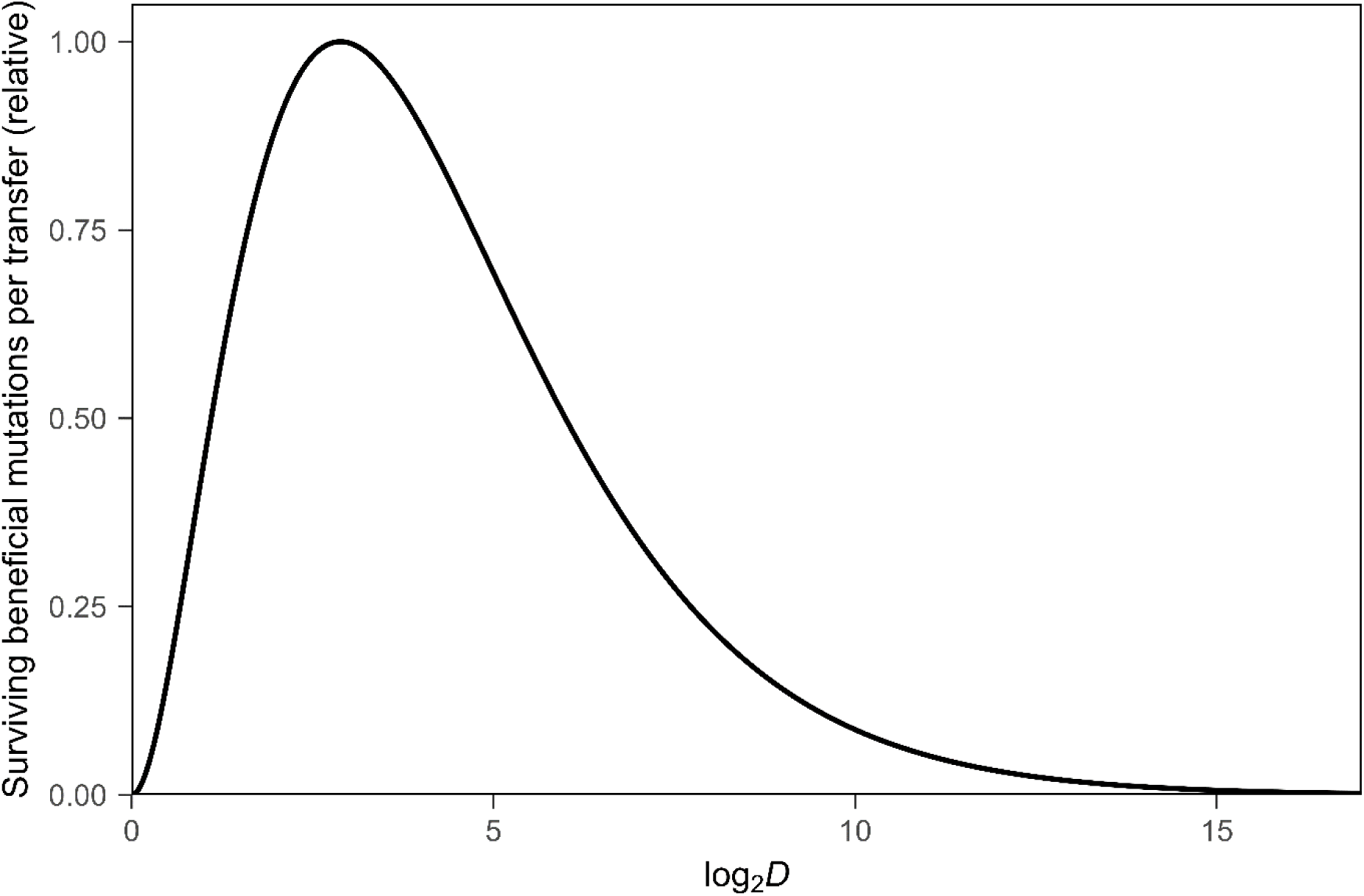
Effect of varying the dilution factor, *D*, on the number of surviving beneficial mutations per transfer (relative scale) in asexual populations propagated via periodic serial transfers. The maximum occurs at *D* = e^2^≅ 7.39 (i.e., log_2_ *D* ≅ 2.89), as shown by Wahl et al. (4).

To address these differences between theoretical models and empirical conditions, we first performed numerical simulations with and without the assumption that beneficial mutations are rare. Our simulations take advantage of the knowledge gained from the LTEE, including previous estimates of the beneficial mutation rate, distribution of fitness effects associated with beneficial mutations, and other relevant parameters (14). The ancestral strain used in the LTEE has a low point-mutation rate, on the order of 10^-10^ per base pair and ∼10^-3^ across the entire genome (13, 32). Both the theoretical model of Wahl et al. (4) and our simulations ignore the fixation of deleterious mutations, which is justified by the low genome-wide mutation rate and the large minimum population size (i.e., during the transfer bottleneck).

We also performed an evolution experiment using bacteria to examine the effect of different dilution treatments on the rate of adaptive evolution in the laboratory. To that end, we propagated 12 populations of *E. coli* in each of four daily dilution treatments (2-, 8-, 100-, and 1000-fold) for 150 days. These treatments are such that the minimum population size is >10^5^ cells even with the most severe bottleneck. We used the same ancestral strains and culture medium as the LTEE, and we performed competition assays to measure changes in the bacteria’s competitive fitness. One methodological difference in our experimental setup from the LTEE is that each population began with an equal mixture of two ancestral strains that differ by a neutral genetic marker, whereas each LTEE population began with one or the other variant. By starting each population with a mixture of the two variants, and by monitoring their relative abundance over time, we could determine when a beneficial mutation began to sweep through the population, causing a shift in the ratio of the marked lineages. For the numerical simulations, we assume that nothing else changes about the environment and the growth rates that govern selection when we vary the dilution factor; the model of Wahl et al. (4) makes the same assumption. In our experiments, however, we examine whether this assumption is violated.

Several experimental studies have recently explored the effects of population bottlenecks on bacterial evolution (33–37). However, some of the experiments imposed a single bottleneck rather than repeated bottlenecks, while others varied the selective environment, final population size, or both. Hence, these experiments do not answer the question posed by Wahl et al. (4), and which we examine in our study: What dilution factor maximizes fitness gains in large populations that experience periodic bottlenecks while reaching the same population size between successive bottlenecks?

## Results and Discussion

### Simulations with Rare Beneficial Mutations

Wahl et al. (4) showed that the supply of beneficial mutations that survive periodic bottlenecks is highest when *D* ≈ 7.4. If beneficial mutations limit adaptive evolution, then the same dilution factor should also maximize the rate of fitness increase. However, that dilution factor might not maximize the rate of fitness increase if beneficial mutations are common. Indeed, as Wahl et al. pointed out, when beneficial mutations are common, many that survive drift loss during the bottlenecks will nonetheless go extinct owing to clonal interference between competing beneficial mutations (22). Therefore, the results of Wahl et al. (4) should be most relevant for predicting the rate of adaptive evolution when populations evolve under the SSWM regime (19–21).

To examine these issues, we developed a numerical simulation package, which allows one to vary the dilution factor and rate of beneficial mutations (along with other relevant parameters). Our simulations track the abundance of every lineage with a new beneficial mutation; a lineage can go extinct by random drift during the periodic dilutions, or by being outcompeted by other lineages that have acquired mutations that confer a greater advantage. Like the model of Wahl et al. (4), our simulations assume that the final population size before each dilution event is constant, and that selection acts only through differences in growth rate. To parameterize the simulations, we use the same final population size as in the LTEE along with estimates derived from the LTEE for the ancestral distribution of fitness effects and the strength of diminishing-returns epistasis (14). The fitness effects of beneficial mutations are drawn from an exponential distribution, with the expected benefit declining with increasing fitness (see Methods and Supplemental Text). These parameters were previously shown to describe and even predict with high accuracy the fitness trajectories observed in the LTEE (14, 38) when using a rate of beneficial mutation also estimated from the LTEE. However, in this paper, we will first use a much lower rate of beneficial mutation in order to confirm that the optimal *D* under the SSWM regime agrees with the theory of Wahl et al. Our simulations, like that theory, should account for the supply of beneficial mutations that survive the bottlenecks and the subsequent growth of the surviving mutants.

To that end, we ran simulations with the beneficial mutation rate set to 1.0 × 10^-11^ per genome. With a final population size of 5 × 10^8^ cells, this rate produces, on average, only 0.005 beneficial mutations per transfer cycle (i.e., 1 mutation in 200 cycles). Moreover, most mutations are lost to drift while they are rare, so that surviving beneficial mutations are few and far between. Therefore, we ran 10,000 simulations for 1500 transfers at each dilution factor in order to have enough replicates that experienced some adaptation to allow a meaningful analysis.

Figure 2 shows the grand mean fitness trajectories during 1500 transfers for ten dilution factors ranging from 2-fold to 100,000-fold under this SSWM regime. The populations propagated with *D* = 8 achieved the highest fitness levels, with progressively smaller gains at both lower and higher dilutions (Table S1). The optimal *D* appears to change slightly during the early transfers (Fig. S1). These changes may reflect noise associated with the small number of surviving beneficial mutations and differences in the time required for new beneficial mutations to reach meaningful frequencies (i.e., where they impact average fitness) as a function of the dilution factor. However, when the runs were extended to 100,000 transfers, and thus encompassed many more surviving beneficial mutations, *D* = 8 clearly achieved the highest fitness gains (Fig. S2, Fig. S3). Thus, our simulations support the finding of Wahl et al. (4) that a dilution factor of that magnitude maximizes fitness gains, assuming that evolution occurs in the SSWM regime where beneficial mutations are rare. However, that assumption requires that the beneficial mutation rate be much lower than it is in the LTEE and many other study systems.

**Fig. 2.**
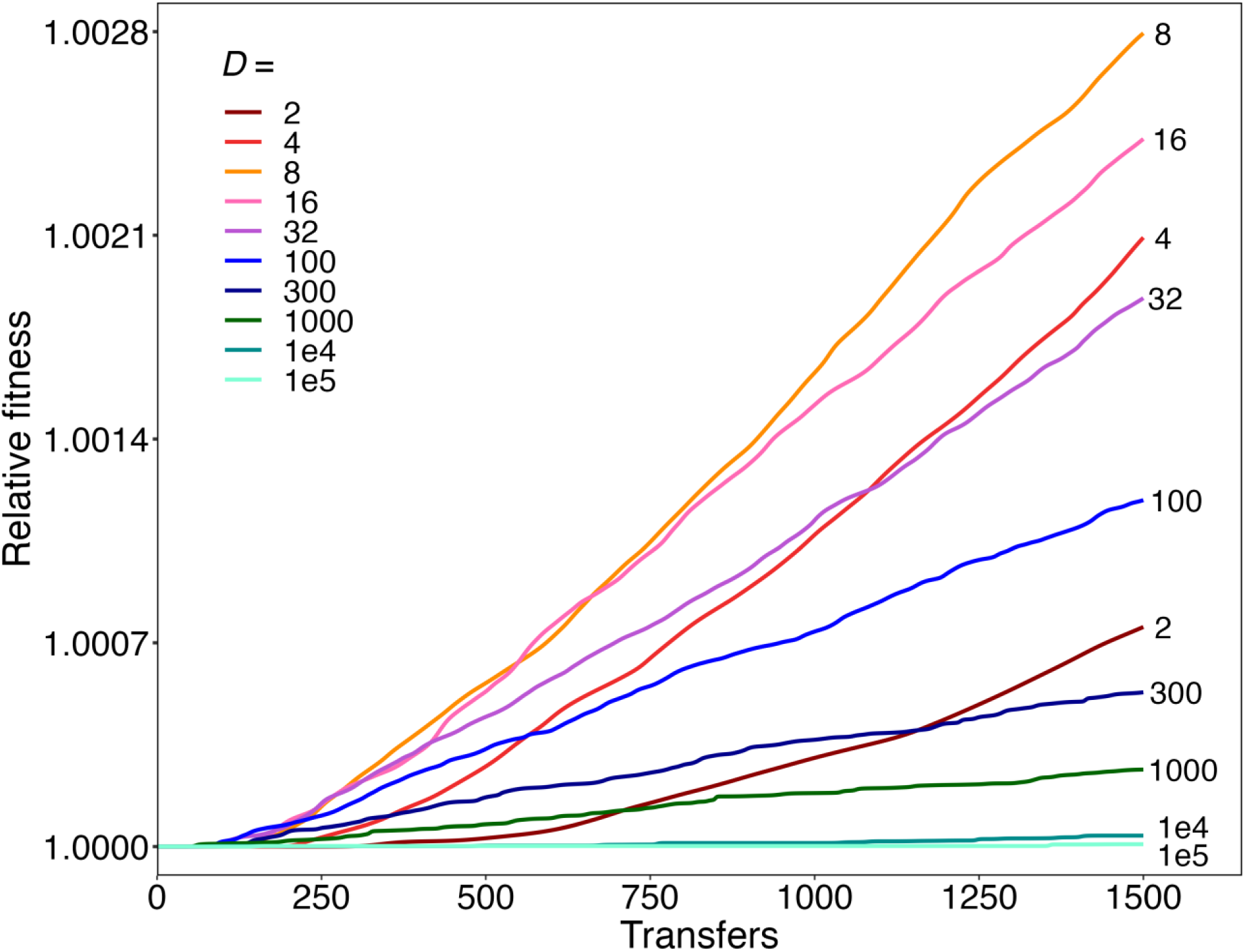
Simulations showing trajectories of relative fitness for populations evolving under a strong selection, weak mutation (SSWM) regime with dilution factors, *D*, ranging from 2-fold to 10^5^-fold. Each trajectory shows the grand mean fitness based on 10,000 runs for 1500 transfer events. Individual runs for *D* = 2, 8, 100, and 1000 are shown in Fig. S4. See text for details.

In this SSWM regime, most populations had no surviving beneficial mutations after 1500 transfers (Table S1). For the populations with fitness gains, each individual trajectory typically has only a single step-like increase, with the height of the step determined by the effect size of the surviving beneficial mutation (Fig. S4). Another revealing feature of the individual trajectories is evident in the slope during the step-like increase in fitness. When we compare trajectories for runs that experienced similar fitness gains, the slope is steeper when the dilution factor is higher than when it is lower (Fig. S4, panels A-D). This difference occurs because the higher dilution factor leads to more generations per transfer cycle, which allows a surviving beneficial mutation to sweep through the population in fewer cycles. The difference in steepness disappears, however, when the fitness trajectories are shown with time expressed as the number of generations (Fig. S4, panels E-H).

### Simulations with Abundant Beneficial Mutations

We then sought to examine the effects of violating the assumption that beneficial mutations are rare on the fitness trajectories and the resulting optimal dilution factor. To that end, we ran simulations using the beneficial mutation rate of 1.7 × 10^-6^ per genome estimated from the LTEE (14), while keeping all other parameters the same. Given this beneficial mutation rate, population size, and distribution of fitness effects, the evolving populations should experience a strong-selection, strong-mutation (SSSM) regime, in which beneficial mutations are both common and have large fitness effects. In this regime, multiple clones with different beneficial mutations are typically present at the same time, and the resulting competition among them will interfere with their progress towards fixation (22–29).

We simulated ten dilution treatments and ran 100 replicates each for 1500 transfers, using the high beneficial mutation rate. Each population had at least one surviving beneficial mutation by transfer 1500 under this SSSM regime; therefore, we did not need to run as many replicates as in the SSWM regime. Figure 3 shows the grand mean fitness trajectories for each dilution factor; Figure S5 shows the trajectories for the replicate populations. These simulations have several noteworthy features. First, the fitness trajectories show a declining rate of improvement over time (Fig. 3), similar to the LTEE (39). The deceleration reflects the impact of diminishing-returns epistasis, such that the fitness effects of beneficial mutations tend to be smaller in more-fit genetic backgrounds (14). In contrast, the mean fitness trajectories in the mutation-limited regime are nearly linear after an initial lag (Fig. 2), even though the diminishing-returns parameter is identical for the SSWM and SSSM simulations. Diminishing-returns epistasis only reduces the fitness gains produced by second and later beneficial mutations, and the populations in the SSWM regime rarely had even one surviving beneficial mutation by transfer 1500 (Table S1). The quasi-linear increase in the grand mean fitness thus results from the independently evolving populations fixing single mutations at different times in the replicates. When the SSWM trajectories are extended to 100,000 transfers, the effect of diminishing-returns epistasis becomes apparent (Fig. S2). However, the curvature of the trajectories is still less pronounced in the SSWM simulations than in the SSSM simulations, where the cumulative fitness gains (and thus the impact of diminishing returns) are much greater (Fig. 3).

**Fig. 3.**
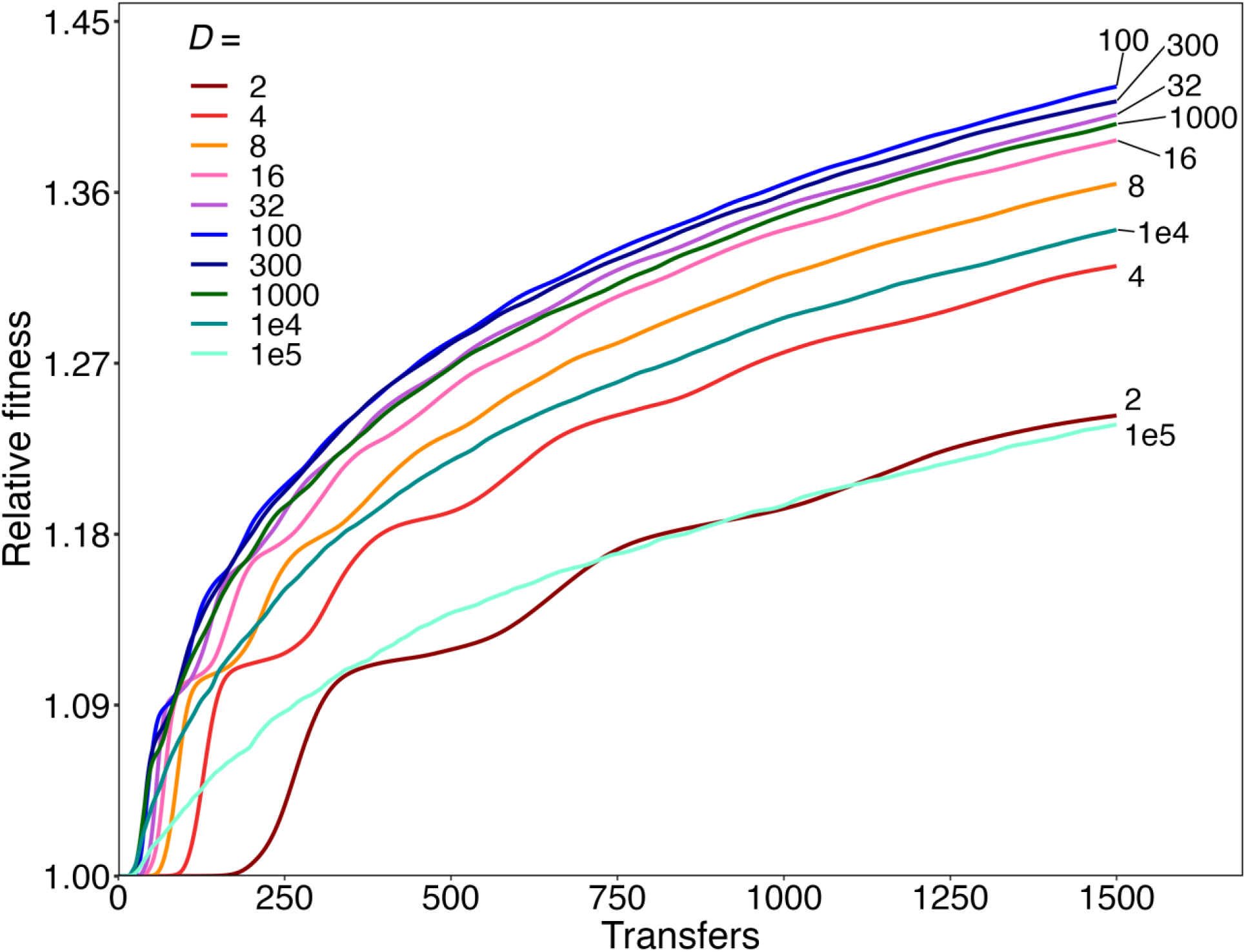
Extended simulations showing trajectories of relative fitness for populations evolving under the strong selection, strong mutation (SSSM) regime with dilution factors, *D*, ranging from 2-fold to 10^5^-fold. Each trajectory shows the grand mean fitness based on 100 runs for 1500 transfer events. Individual runs are shown in Figs. S5 and S7.

Second, there are large step-like increases in the grand mean fitness trajectories under the SSSM regime when *D* is small (Fig. 3). These steps reflect an initial lag period that depends on the time required for the first selective sweep to become evident in a typical population (Fig. S5). These steps are more noticeable when *D* is small because each transfer cycle allows fewer generations and thus beneficial mutations require more transfers to sweep to fixation (Fig. S6). By contrast, when one examines the fitness trajectories for individual populations in these simulations, they all show step-like increases in fitness (Fig. S5, Fig. S7). Indeed, step-like fitness dynamics are an expected feature of evolving asexual populations (3, 22, 39). The size of each step reflects the benefit conferred by a mutation (or sometimes multiple linked mutations) sweeping through the population.

Third, and most importantly, by 150 transfers, the highest mean fitness was achieved with *D* = 100 in the SSSM regime, followed by dilution factors of 300 and 32 (Fig. S6, Fig. S8). Table S2 shows that the final mean fitness values differ significantly for all pairs of adjacent dilution factors, even though the magnitude of the difference is small in some cases (e.g., *D* = 100 versus *D* = 300). Thus, while our simulations confirmed the prediction that ∼8-fold dilutions are optimal for maximizing fitness gains in the mutation-limited SSWM regime (Fig. 2), they also demonstrate that 8-fold dilutions are far from optimal when beneficial mutations are common and clonal interference is important. Notice also that the mean fitness trajectories consistently fall away at both higher and lower dilution factors, which highlights the general tradeoff. On the one hand, a lower dilution factor allows more beneficial mutations to survive the transfer bottlenecks. On the other hand, higher dilutions result in more generations between the successive transfers, allowing the fitness advantage of beneficial mutations to compound at a faster rate on a per-transfer basis. Moreover, when beneficial mutations are common, multiple lineages that harbor independently arising beneficial mutations must compete with one another, which evidently increases the dilution factor that maximizes the rate of adaptation.

It is also noteworthy that the grand mean fitness trajectories appear to become strikingly parallel across a wide range of dilution factors, after the more complex trajectories during the early transfers (Fig. 3). To verify this impression, we calculated the change in fitness over the last 500 transfers for each replicate population (Fig. S5), and we compared the average change between adjacent dilutions (Table S3). There were no significant differences in the fitness gains over these final transfers. In fact, the fitness gains during this period were similar even when the dilution factor ranged from 2-fold to 10^5^-fold.

These simulations demonstrate that ∼8-fold dilutions do not produce the greatest fitness gains in the SSSM regime, where beneficial mutations are common. In fact, there is no single dilution factor that maximizes gains for all other parameter values in this regime. To demonstrate this fact, we ran simulations where we changed the mean effect size of beneficial mutations to values that were lower or higher than estimated for the LTEE while keeping all other parameters the same (Fig. S9). Dilution factors of 100, 300, and 1000 produced similar grand mean fitness trajectories when the initial beneficial effect size was reduced to one-third or even one-tenth of the estimate based on the LTEE. When the initial effect size was increased three-fold, dilution factors of 16, 32, and 100 produced the greatest fitness gains; and with a ten-fold increase in the beneficial effect size, dilution factors of 8, 16, and 32 had the greatest gains.

### Marker Divergence in Evolution Experiment with Bacteria

Our simulations show that SSWM and SSSM regimes—with beneficial mutations being rare and abundant, respectively— produce fitness trajectories with distinctive features. Moreover, the dilution factors predicted to maximize fitness gains differ substantially between these regimes: ∼8-fold dilutions are optimal in the SSWM regime, whereas ∼100-fold dilutions are optimal in most of the SSSM regimes that we tested. To examine the effects of the dilution factor on the dynamics of adaptation in an actual biological system, we performed an evolution experiment using the same ancestral *E. coli* strains and culture medium as in the LTEE (see Materials and Methods), except with four dilution treatments. To that end, we propagated 48 populations for 150 days with 2-, 8-, 100-, or 1000-fold daily dilutions of the previous day’s culture in fresh medium. Each day, a population grew for log_2_ *D* cell generations until it reached its final population size, at which point the limiting glucose was depleted. The four treatments thus allowed 1, 3, ∼6.6, and ∼10 generations per daily transfer, respectively, which correspond to 150, 450, 1000, and 1500 generations over the full experiment.

All populations started with an equal mix of Ara^−^ and Ara^+^ strains that have equal fitness in the glucose-limited medium but differ by a mutation that causes cells to form red and white colonies, respectively, when plated on tetrazolium arabinose (TA) agar (3). The ratio of the two lineages should remain constant over time, except for statistical fluctuations (primarily sampling error when only a few hundred colonies are counted), until a beneficial mutation arises and spreads through one or the other lineage. Beneficial mutations cause sustained perturbations to the ratio in a given population, but across multiple populations the direction of the perturbation is random. Comparing the trajectories of the ratio between treatments allows us to quantify differences in the timing of the perturbations caused by beneficial mutations. Each marked lineage within a particular treatment started from an inoculum obtained by plating for single colonies (each derived from a single cell) to ensure that the replicate populations did not share any mutations that were identical by descent (40). However, the same two source colonies were used as founders for one population in each treatment. We sampled each evolving population every 3 days and spread the sample on a TA agar plate, allowing us to score cells based on whether they were descended from the Ara^−^ or Ara^+^ founder. We continued this procedure until a population appeared to have fixed one or the other marker state, after which we plated samples every 15 days.

Figure 4 shows the resulting trajectories of the log_2_-transformed ratio of the abundances of the marked lineages for the 12 populations in each dilution treatment. For 10 populations in each treatment, the ratio remained essentially constant for several weeks, as expected given the marker’s neutrality. However, for 2 populations in each treatment, the marker ratio systematically increased (grey trajectories) or decreased (mauve trajectories) from the beginning of the experiment. These deviations imply that the founding clones already harbored mutations that were not neutral. This inference is supported by the fact that the same replicates (founded from the same pair of colonies) experienced the same directional deviations in all four treatments. Having saved frozen samples of the 24 founding colonies, we tested whether those particular clonal isolates had mutations that affected their fitness by competing them against the reciprocally marked LTEE ancestors using the 100-fold dilution protocol. Indeed, the founding clones from the two atypical populations differed significantly in fitness (Table S4), and in the directions expected from their aberrant trajectories (Fig. 4), confirming the presence of non-neutral mutations in the founders. We therefore excluded these two populations in all four dilution treatments from our subsequent analyses, because they violate the assumption of initial neutrality. The trajectories of the 10 remaining populations suggest that deviations from the initial 1:1 ratio (0 on the log-transformed scale) of the marked lineages tended to start earlier in the populations that were subject to the more severe bottlenecks (Fig. 4).

**Fig. 4.**
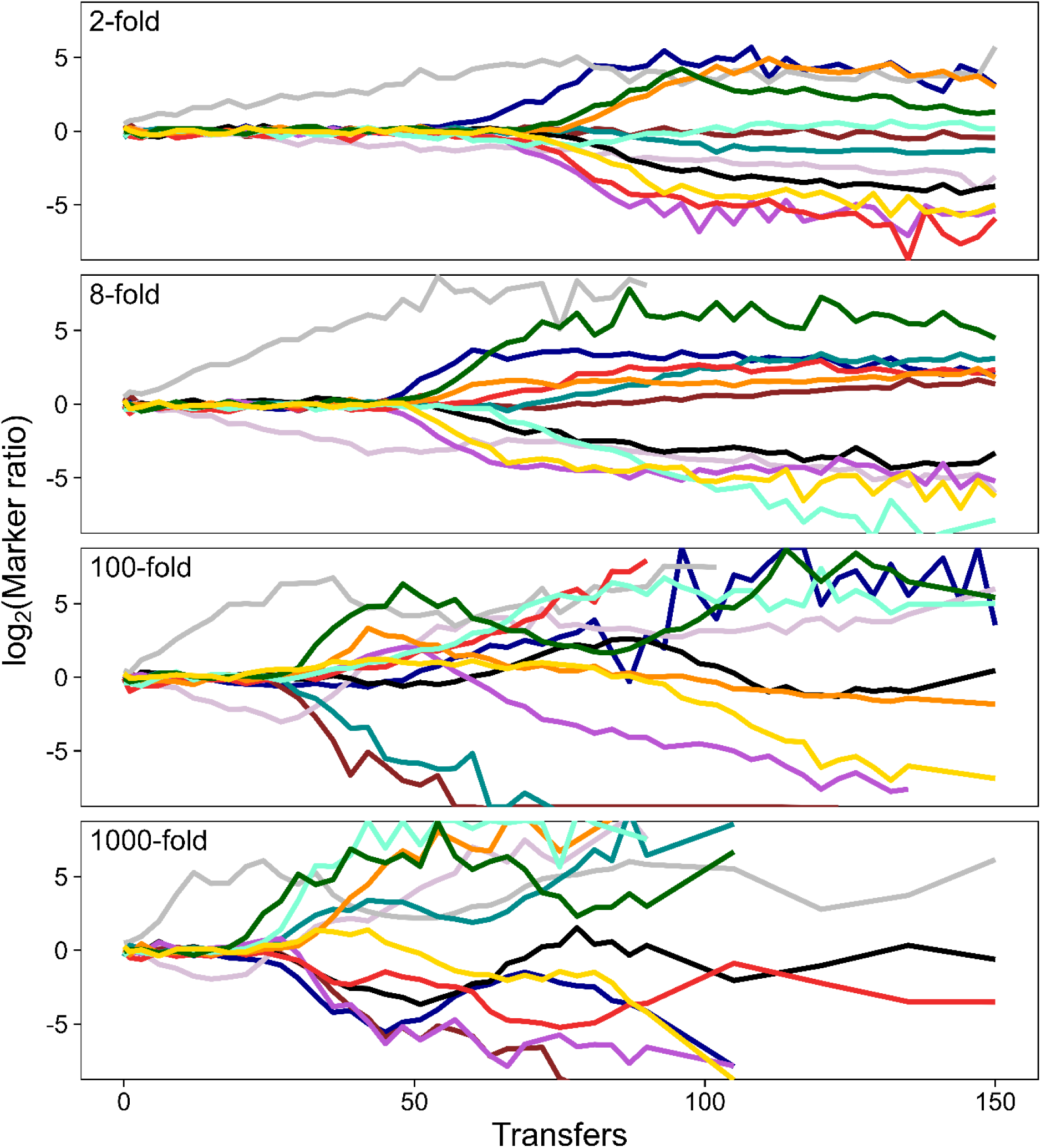
Trajectories of the relative abundance of two marked lineages in 12 *E. coli* populations evolving in each of 4 dilution treatments for 150 daily transfers. With a neutral marker, the ratio remains essentially constant until a beneficial mutation sweeps through one of the marked lineages. Two populations (gray and mauve lines) in each regime were founded by the same pair of clones that had inadvertently acquired non-neutral mutations; they were excluded from all subsequent analyses. Note the log_2_-transformed scale for the marker ratio.

### Time to Divergence in the Experiment and Simulations

To confirm that the marker-ratio trajectories started diverging significantly earlier in the experimental populations that experienced more severe bottlenecks, we calculated the time to divergence, *TD*, for each population, based on when the ratio diverged from 1:1 by at least a factor of 2 (i.e., the log_2_ ratio deviated from 0 by 1) in either direction. One population in the 2-fold dilution treatment never deviated by 2-fold from the 1:1 ratio, and we used 150 days (i.e., the duration of the experiment) as its *TD* for this analysis.

Figure 5 shows the *TD* values obtained for the experimental populations in all four dilution treatments. All six pairwise comparisons between the treatments are significant (Table S5), even after a Bonferroni correction for multiple comparisons. Moreover, a log-log regression of *TD* on *D* is highly significant (*t* = –10.63, 38 df, *p* < 0.0001). Thus, the marker ratios diverged earlier with increasing values of *D*, indicating that selective sweeps began earlier at the higher dilutions.

**Fig. 5.**
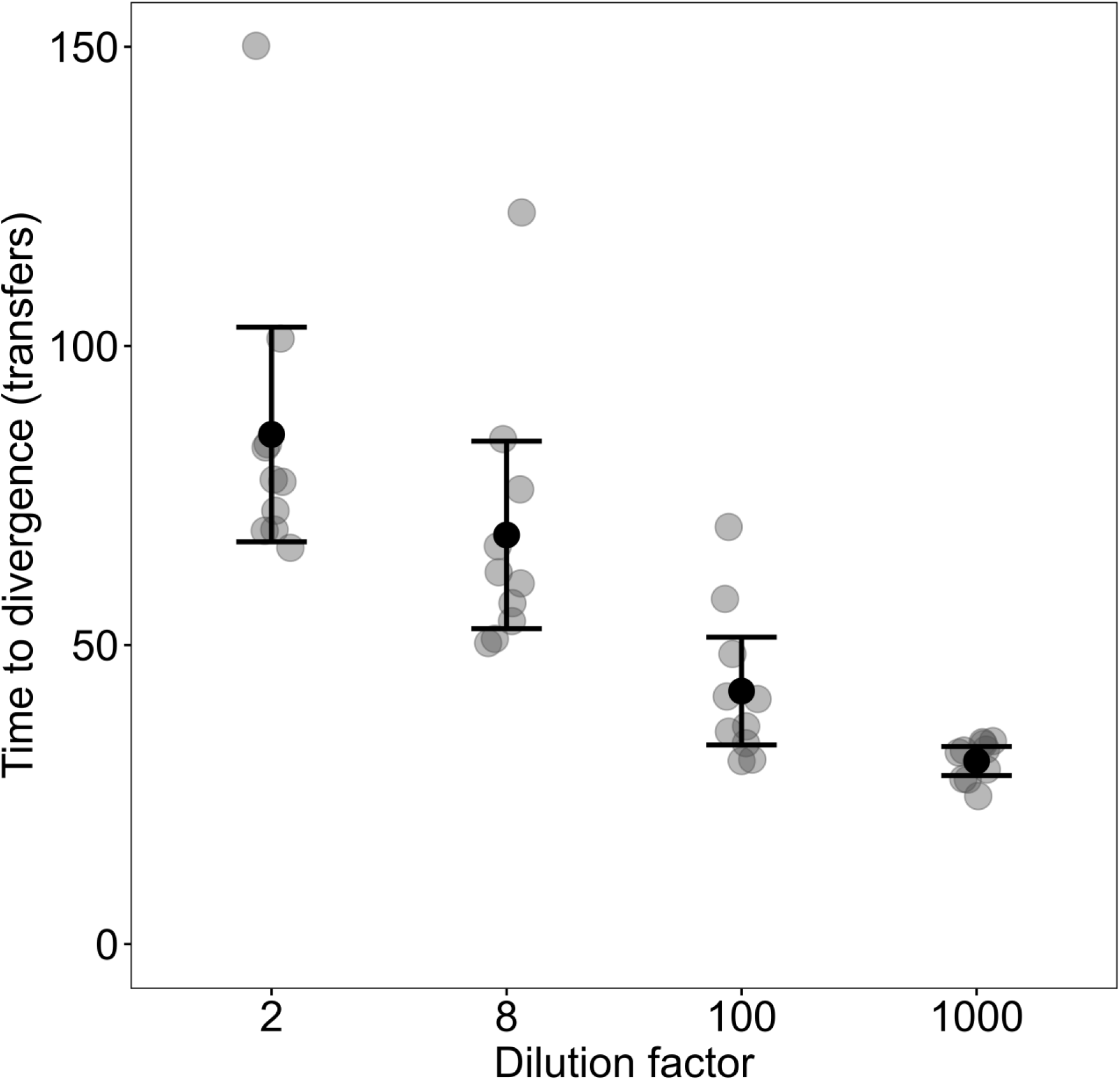
Time to divergence, *TD*, of the marker ratio in populations propagated under different dilution treatments. For each population, *TD* is defined as the time of the first sample when the ratio of the neutrally marked lineages deviated from unity by two-fold in either direction, based on counts made every third day. Black and gray symbols show means and replicate assays, respectively; error bars are 95% confidence intervals. See text for details and Table S5 for statistical analysis.

To better understand these results, we also analyzed the marker-ratio trajectories for the simulations under the SSSM regime across the extended range of dilution factors (Table S6). These simulations show earlier marker divergence times for 300-to 10^4^-fold dilutions than for lower or higher dilution factors. The simulated trajectories thus indicate that the average time to divergence is minimized at an intermediate dilution factor. However, the dilution factor that minimizes the time to divergence (300 < *D* < 10^4^) differs from the one that maximizes long-term fitness gains (32 < *D* < 300; Table S2). For the four dilution factors used in both the simulations and experiment, the divergence times show similar trends, with one conspicuous exception: namely, for *D* = 2 the marker ratio took much longer to diverge in the simulations than in the experiment. In fact, none of 100 simulations at that dilution factor showed divergence within 150 transfers, whereas 9 of the 10 experimental populations had diverged by that timepoint. We will see further evidence of this discrepancy for *D* = 2 when we examine the changes in fitness in the experimental populations.

### Fluctuations in the Marker Ratio

We also examined the magnitude of the fluctuations in the marker ratio between consecutive experimental samples through day 21 (just before the earliest 2-fold divergence in any of the 40 qualifying populations). We did so to check whether the dilution regimes produced different fluctuation patterns that might confound our use of the same criterion (i.e., a 2-fold shift in ratio of the marked lineages) to identify perturbations caused by selective sweeps of beneficial mutations. We did not expect to see different fluctuation profiles across the treatments, despite the different bottleneck sizes, because the sampling error associated with scoring a few hundred cells in a typical sample is much greater than the effect of random drift in populations with >10^5^ cells even during the most severe experimental bottleneck treatment (i.e., ∼5 × 10^7^ cells/mL × 10 mL, then diluted 1000-fold). As expected, an ANOVA shows no significant differences among the treatments (Table S7). Also, only 3.6% (10/280) of the fluctuations were even half as large as the 2-fold change in marker ratio used as the cutoff to identify perturbations caused by beneficial mutations. Moreover, those 2-fold changes were generally followed by further large changes (Fig. 4), as expected for selective sweeps, but not for fluctuations caused by drift or sampling noise.

Readers who are unfamiliar with marker-divergence experiments may, on first inspection, be surprised by the reversals in the marker-ratio trajectories that occurred in some populations after the onset of a selective sweep (Fig. 4). However, such reversals are widely observed, and indeed expected, given clonal interference between contending lineages that have independently acquired beneficial mutations (22, 26, 41, 42). As a beneficial mutation sweeps through one marked lineage, it must compete not only against the progenitors (of both marker types) but also against any beneficial mutations that arose in the opposing lineage. If the competing mutations have similar fitness effects, then the marked lineages might appear to equilibrate at some new ratio. If the later-arising mutation is more beneficial than the one that arose first, then the marker-trajectory may reverse direction, and further reversals can occur as additional beneficial mutations arise in one or both lineages.

### Fitness in Evolved and Common Environments

To quantify the extent of adaptation in the experimental populations, we measured the fitness gains of the final evolved bacteria (from day 150 of our experiment) by competing them against the ancestral strain. We could not measure the fitness of the whole-population samples because we had to mix the evolved bacteria with the ancestral strain bearing the alternative marker to measure their relative fitness, and some of the populations still had cells from both marked lineages at the end of the experiment (see Materials and Methods). Instead, we isolated clones from each endpoint population, and we competed them against the reciprocally marked ancestor.

Before comparing the fitness of the strains evolved in the different dilution treatments, we first sought to determine whether adaptation is interchangeable across the treatments. An implicit assumption of both the theory and simulations is that fitness is simply the growth rate of a mutant carrying a beneficial mutation relative to that of its parent. In reality, other demographic (18) or ecological (43) factors could come into play. For example, fitness might be more sensitive to changes in the duration of the lag phase at lower dilutions, and the effect of metabolic byproducts carried over in the medium during transfers might be more important at lower dilutions. To address this issue, we measured the fitness of each evolved clone in two conditions, which we call the “evolved” and “common” environments. For the former, we measured fitness using the same dilution protocol in which the clone had evolved. For the latter, we used the 100-fold dilution protocol, regardless of the treatment in which the clone evolved.

Figure 6 shows the relative fitness values for the 150-day clones (excluding the two clones from the aberrant populations in each treatment), plotted as the values in their evolved environment versus in the common environment. The clones that evolved in the 2-fold dilution treatment were significantly more fit in their evolved environment than the common environment (Fig. 6A, Table S8). They had an average fitness gain of ∼24% with 2-fold dilutions, but only ∼6% when subjected to 100-fold dilutions. Thus, the nature of selection must have differed between the 2-fold and 100-fold treatments. This difference might reflect the heightened importance of shortening the lag phase relative to faster growth when there is only one doubling per day, or the fact that half of the medium contains potential byproducts from the previous day with a 2-fold dilution (as opposed to only 1% carryover with a 100-fold dilution).

**Fig. 6.**
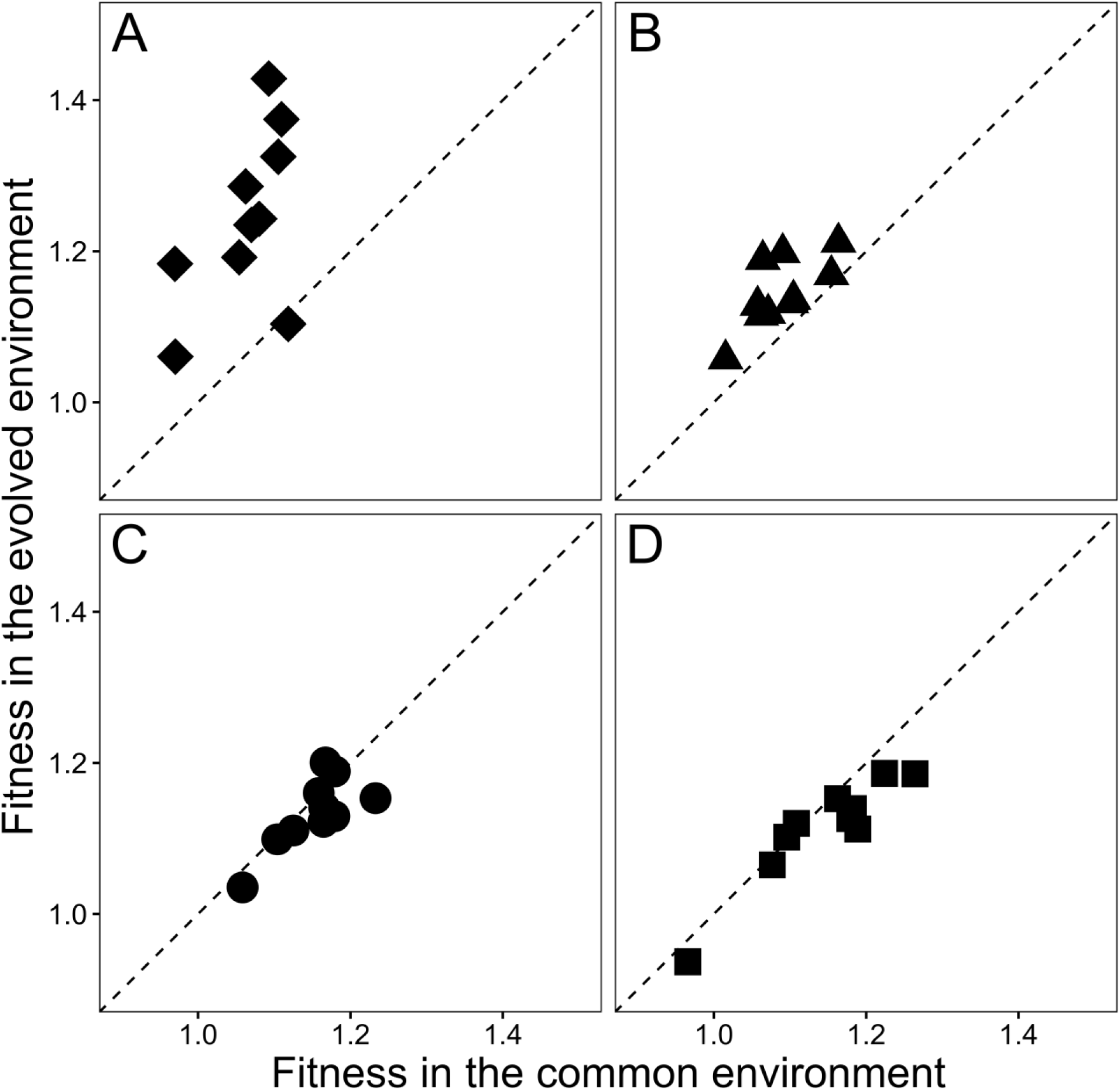
Comparisons of fitness measured in the evolved and common environments for bacteria propagated by 2-fold (A), 8-fold (B), 100-fold (C), and 1000-fold (D) dilutions for 150 days. In the evolved environment, bacteria sampled at day 150 competed against a marked ancestor in the same dilution treatment where they had evolved. In the common environment, competition assays were performed using 100-fold dilutions, regardless of the evolutionary treatment. Dashed lines show the expectation if fitness values are equal in the two environments.

The clones that evolved in the 8-fold dilution treatment also tended to be more fit in their evolved environment than in the common environment with 100-fold dilutions, but this difference was much smaller than for the 2-fold treatment (Fig. 6B, Table S8). To see whether the difference in fitness between the evolved and common environments of the clones in the 8-fold treatment might undermine using the common environment for comparison purposes, we measured the fitness of the clones from both the 8-and 100-fold treatments under the 8-fold dilution protocol (Fig. S10A). The clones that evolved in the 100-fold treatment had significantly higher fitness than those that evolved in the 8-fold treatment, even when the competitions were performed with the 8-fold protocol (*p* = 0.0195, two-tailed Wilcoxon’s signed-ranks test). Also, the values measured using the 8-fold and 100-fold dilutions were well correlated for the clones from both evolutionary treatments (Fig. S10B, *r* = 0.8218, *p* < 0.0001). Thus, selection in these treatments was sufficiently similar that evolution in the 100-fold dilution treatment led to higher fitness than evolution in the 8-fold treatment, regardless of the competition environment.

Therefore, the fitness gains measured in the evolved and common environments for the bacteria that evolved in the 8-, 100-, and 1000-fold dilution treatments are largely consistent with simple selection for faster growth. However, the bacteria that evolved in the 2-fold treatment experienced substantially different selection pressures. Thus, that treatment cannot be used to test predictions about the dilution factor that maximizes fitness gains based on the theoretical model and numerical simulations, both of which assume equivalent selection for faster growth across all dilutions.

### Comparison of Fitness Gains in Bacteria that Evolved in Different Treatments

We ran additional competitions with 5-fold replication to obtain more accurate estimates of the fitness levels of the clones that evolved for 150 transfers in the 8-, 100-, and 1000-fold dilution treatments, when all were measured using the common 100-fold protocol (Fig. 7). The populations from the 100-and 1000-fold treatments were significantly more fit than those from the 8-fold treatment (Fig. 7, Table S9), whereas there was no significant difference between the 100-and 1000-fold treatments. These direct measurements of fitness therefore support the findings from the SSSM simulations of faster adaptation with 100-and 1000-fold dilutions than with 8-fold dilutions, and they are contrary to the model and simulations that assume a mutation-limited SSWM regime.

**Fig. 7.**
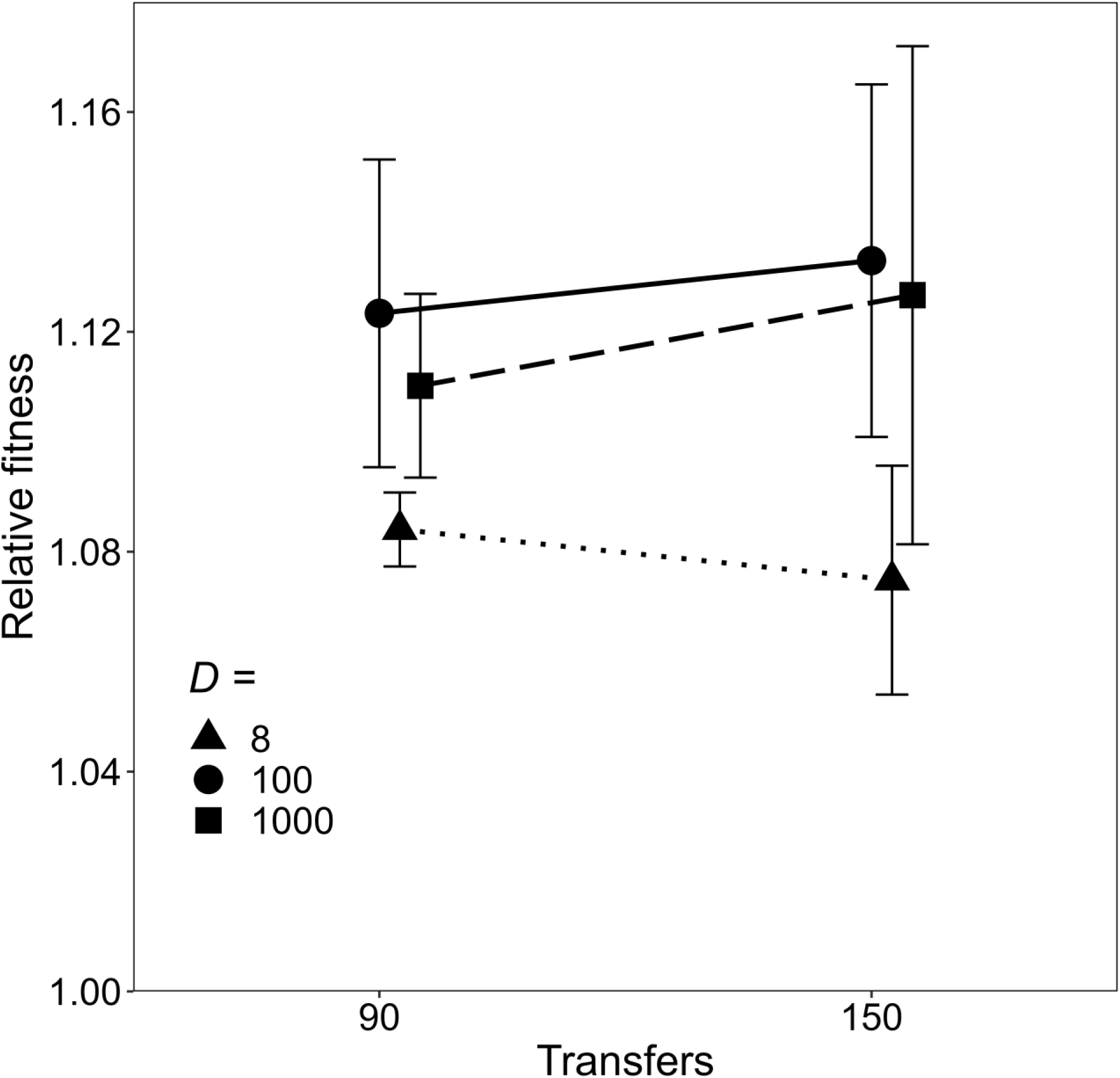
Relative fitness measured after 90 and 150 transfers for bacteria that evolved in the 8-fold, 100-fold, and 1000-fold dilution treatments. Error bars show 95% confidence intervals based on 10 replicate populations in each treatment.

### Fitness Improvement during Later Transfers

Our experiment shows that selective sweeps began earlier (Fig. 4, Fig. 5) and cumulative fitness gains were larger (Fig. 7, Table S9) with *D* = 100 and *D* = 1000 than with *D* = 8. Our simulations in the SSSM regime show similar outcomes (Fig. 3, Fig. S6, Table S6). By contrast, the theoretical model and simulations under the SSWM regime predict that adaptation should be maximized when *D* ≍ 8. Thus, it seems that the assumption of whether beneficial mutations are common or rare can explain these different outcomes. Before accepting that explanation, however, we also considered the following possible complication. The model implicitly assumes a steady-state scenario, whereas both simulated and actual populations deviate from steady-state dynamics. In particular, evolving asexual populations generally exhibit time lags before their fitness trajectories begin to increase appreciably because they begin without any genetic variation for fitness.

To address this potential complication, we examined the change in mean fitness between 90 and 150 transfers in the experimental populations (Fig. 7). We chose 90 transfers as the earlier timepoint because the ratio of the marked lineages had diverged, and thus measurable adaptation was underway, in the three relevant treatments by that timepoint (Fig. 4). As before, we excluded the populations that harbored the non-neutral mutations that caused their marker ratios to deviate from the outset of the experiment. The average fitness increases over these 60 transfers were higher for both the 100-and 1000-fold dilution treatments than for the 8-fold treatment (Fig. 7), although the differences were not statistically significant (Table S10). The slight decline in the average fitness in the 8-fold treatment is not significant; it presumably reflects measurement error, which is unavoidable in experimental data. In any case, there is no evidence that the fitness improvement in the 8-fold dilution treatment would, after a slow start, overtake the gains observed in the 100-and 1000-fold treatments.

### Summary of Key Findings and Conclusions

We performed simulations and experiments to understand how varying the dilution factor in serial-transfer protocols affects the dynamics of adaptation in large populations of asexual organisms. In particular, we examined the scenario in which both the final population size reached in each transfer cycle and the number of transfers that take place are held constant. We examined only those cases in which the minimum population size during the transfer bottleneck was large, with at least thousands of cells in our simulations and hundreds of thousands of cells in our experiments. We did so because our focus in this study is on the effects of moderate bottlenecks on the dynamics of adaptive evolution. These effects are not as well understood as those that occur during extreme bottlenecks, which cause random mutations to accumulate with a concomitant loss of fitness over time (11–13).

A theoretical study by Wahl et al. (4) derived the value of the periodic dilution factor, *D*, that maximizes the rate of adaptive evolution under this same scenario. They demonstrated that adaptation should be fastest at *D* ≈ 7.4 (Fig. 1). A key assumption of their analysis is that beneficial mutations that survive the periodic bottlenecks are so rare that the mutations fix individually and sequentially in an evolving population. However, numerous experiments with microbes have shown that beneficial mutations are common in large populations, such that many of them survive moderate bottlenecks. In that case, concurrent lineages that possess different beneficial mutations compete with one another, leading to the extinction of most lineages via clonal interference (22–28).

When we performed numerical simulations of evolving populations across a wide range of dilutions—with parameters estimated from the canonical *E. coli* LTEE, except using a much lower rate of beneficial mutation—we confirmed the theoretical prediction that ∼8-fold dilutions produce the highest fitness gains (Fig. 2). In these simulations, however, only a minority of the populations in any treatment had substituted even a single beneficial mutation after 1500 transfers (Table S1). With daily propagation in a laboratory, 1500 transfers would require over 4 years, and still most populations would not have undergone any adaptation whatsoever.

Using the same parameters, but with the higher rate of beneficial mutations estimated in the LTEE, adaptation was evident within just 150 transfers across a wide range of dilution factors. Among the ten treatments in our simulations, we saw the highest average fitness gains after 150 transfers with 100-and 300-fold dilutions (Fig. S6). By 1500 transfers, the 100-fold treatment produced the maximum gains by a small, but significant, margin (Fig. 3, Table S2).

In both cases, therefore, fitness gains are maximized at an intermediate dilution factor. An intermediate optimum occurs when beneficial mutations are rare, as shown for the SSWM regime analyzed by Wahl et al. (4) and seen in our simulations with a low beneficial mutation rate (Fig. 2). An intermediate optimal value for the dilution factor also exists when beneficial mutations are common, as shown by our simulations with a high rate of beneficial mutations (Fig. 3). However, the optimal dilution factor shifted to a higher value as competition between multiple lineages with different beneficial mutations became important. In this case, the specific location of the optimum value also depended somewhat on whether the optimum dilution factor was defined by the average time required for the earliest beneficial mutations to sweep through evolving populations (Table S6) or by the average long-term fitness gains (Fig. 3).

On balance, the results of our experiment with *E. coli* populations were consistent with the simulations that used the high rate of beneficial mutation estimated from the LTEE. However, the experiment had lower resolution than the simulations for several reasons: fewer dilution treatments (4 versus 10); fewer populations per treatment (10 versus 100); shorter duration (150 versus 1500 transfers); and less precise fitness estimates (owing to measurement noise). Even so, measurable adaptation began sooner (Fig. 5) and fitness gains were larger (Fig. 7) in the 100-and 1000-fold dilution treatments than in the 8-fold treatment, consistent with the simulations.

There was, however, one noteworthy discrepancy between the experiment and simulations: namely, the experimental populations subjected to 2-fold dilutions evolved increased fitness earlier (Fig. 4) and to a greater extent (Fig. 6) than did the corresponding simulated populations (Fig. 3, Fig. S6). The simulations assume that genotypes vary only in their exponential growth rates, such that the fitness of any evolved genotype relative to the ancestor (expressed as the ratio of the growth rates) is the same when measured in competitions at different dilution factors. However, the bacteria that evolved in the 2-fold treatment had much higher fitness when competed against the ancestor using 2-fold dilutions than when using 100-fold dilutions (Fig. 6A). Thus, the assumption that fitness gains involved only changes in the exponential growth rate was violated when the bacteria evolved in the 2-fold dilution treatment. By contrast, our results are largely consistent with the assumption that selection favored faster exponential growth for the bacteria that evolved at higher dilutions (Fig. 6).

### Future Directions

Our results show that the effect of periodic bottlenecks on the dynamics of adaptation in large asexual populations involves a tradeoff between the loss of genetic diversity, which is exacerbated by more severe bottlenecks, and the rate of spread of surviving beneficial mutations, which is accelerated by having more generations between severe bottlenecks. However, a number of interesting questions remain for future work. In terms of our experiment with bacteria, we would like to understand why the effects of selection were so different in the 2-fold dilution treatment from the other treatments. With the 8-, 100-, and 1000-fold dilutions, our results were largely consistent with simple selection for faster growth. In the 2-fold regime, selection evidently favored other fitness-related traits. For example, the bacteria might have evolved a shorter lag phase before starting to grow after each transfer (18); or they may tolerate and even exploit secreted metabolites (43), which could be especially important in an environment where half the medium is carried over with each transfer. Whole-genome sequencing of the evolved bacteria might reveal how the genetic targets of selection differ between the 2-fold and other dilution treatments (44), which in turn could provide insights into the phenotypic traits under selection.

In terms of theory, numerical simulations can guide future efforts to identify the dilution factor that maximizes adaptation under a broader set of conditions than those considered by Wahl et al. (4) or in our own study. Further development of a more general theory might enable one to examine the sensitivity of the optimal dilution factor and the resulting rate of adaptation to other potentially relevant parameters, including the rate of deleterious mutations. Like the theory of Wahl et al., we ignored deleterious mutations in our simulations. We did so because the mutation rate in *E. coli* with functional DNA repair is extremely low. From the rate of accumulation of synonymous mutations in the LTEE, the point-mutation rate has been estimated as ∼10^-10^ per bp per generation (32). With a genome of ∼5 × 10^6^ bp, this rate implies <0.001 mutations per genome replication, and many of them will be neutral rather than deleterious. The rates of other types of mutations, including insertions and deletions, are less certain, but mutation-accumulation experiments show that the genome-wide rate of all deleterious mutations is very low (11, 13). However, six of the twelve LTEE lines evolved hypermutability that increased their point-mutation rates by ∼100-fold (13, 15, 45), although some of them later evolved compensatory changes that reduced the elevated rate (13, 15, 46). Moreover, as the LTEE goes forward, the rate of adaptive evolution should continue to slow owing to pervasive diminishing-returns epistasis (14). As a consequence of higher mutation rates and reduced effect-sizes of beneficial mutations, deleterious mutations may become important for understanding the dynamics of adaptive evolution in the LTEE. More generally, deleterious mutations are undoubtedly important in some other evolution experiments, such as with highly mutable RNA viruses or when imposing extreme bottlenecks that allow some deleterious mutations to fix. Therefore, it will be interesting to perform simulations and extend theory (47) to investigate the combined effects of deleterious and beneficial mutations on the dynamics of adaptive evolution in large asexual populations that harbor lineages with multiple beneficial and deleterious mutations, and which are subject to periodic bottlenecks. For example, such work might be used to predict the dilution factor that balances adaptive and maladaptive evolution. Mean fitness increases in large populations that experience moderate bottlenecks, as seen in the LTEE, but mean fitness tends to decline in populations subject to extreme (e.g., single cell) bottlenecks, as in mutation-accumulation experiments (11, 13). Thus, there presumably exists some intermediate bottleneck size where fitness gains and losses tend to offset one another, albeit with stochastic fluctuations in any given population. Identifying and then experimentally probing that quasi-equilibrium are worthwhile goals for future research.

## Materials and Methods

### Computer Simulations

We wrote a program called STEPS—for Serially Transferred Evolving Populations Simulation—to simulate mean fitness and marker-ratio trajectories. The program tracks unique lineages, each defined by one or more mutations, in a population propagated by serial transfers. The STEPS program takes the following parameters as inputs: final population size after growth has exhausted the limiting resource (*N_f_*), dilution factor (*D*), beneficial mutation rate (*μ_B_*), mean effect size of beneficial mutations in the ancestor (*s*_0_), and strength of diminishing-returns epistasis (*g*). Each population starts with an initial size of *N_f_* / *D* that is split between two equally abundant founding lineages that differ only by a neutral marker. Beneficial mutations are introduced at random as the population grows, thereby generating new lineages that are tracked over time. The size of each benefit is drawn from an exponential distribution, with the expected benefit declining as a lineage’s fitness increases. The lineages grow exponentially according to their relative fitness values. Growth continues until the total population reaches its final size. At that point, the population’s mean fitness is recorded along with the ratio of the two markers. The population is then bottlenecked by the specified dilution factor, with survivors determined by randomly sampling the lineages. In all simulations, we used *N*_f_ = 5.0 × 10^8^, as estimated by Lenski et al. (3) for the ancestral strain in the LTEE environment. For simulations in the strong-selection, strong-mutation (SSSM) regime, we used the following parameters estimated from the LTEE by Wiser et al. (14): *μ*_B_ = 1.7 × 10^−6^, *s*_0_ = 0.012, and *g* = 6.0. We used the same parameters for the strong-selection, weak-mutation (SSWM) regime except with *μ*_B_ = 1.0 × 10^−11^. We provide further details regarding implementation of the simulations in the Supplemental Text.

### Evolution Experiment and Dilution Treatments

We isolated 12 clones as single colonies from freezer stocks of each of two *E. coli* strains, REL606 (Ara^−^) and REL607 (Ara^+^). The clones were incubated in Luria-Bertani broth for 24 h in a shaking incubator at 37°C and 120 rpm, then diluted 10,000-fold in Davis minimal medium supplemented with 25 μg/mL glucose (DM25) and incubated for another 24 h in the same conditions. On day 0 of the evolution experiment, we mixed one Ara^−^ isolate and one Ara^+^ isolate equally to make 12 cultures. These mixed cultures were transferred to test-tubes with fresh DM25 medium using four dilution treatments. For the 2-fold treatment, we transferred 5 mL of each mixed culture into 5 mL of DM25. For the 8-fold treatment, we transferred 1.25 mL of each culture into 8.75 mL of DM25. For the 100-fold treatment, we transferred 0.1 mL into 9.9 mL of DM25. For the 1000-fold treatment, we diluted a portion of each mixed culture 10-fold in saline solution and then transferred 0.1 mL into 9.9 mL of DM25. By using the same culture volume and resource concentration, the bacteria reached the same final population size in all four treatments. However, the number of generations (doublings) varied with the treatment, such that the bacteria underwent 1, 3, ∼6.6, or ∼10 generations per daily transfer cycle. We incubated the 48 populations for 24 h in a standing incubator at 37°C and transferred them daily following the same dilution protocols for 150 days. Every 15 days, we froze samples of each population at –80 °C with glycerol as a cryoprotectant (13.3% v/v final concentration).

### Marker Divergence Analysis

During the evolution experiment, we diluted and spread cells from each population on tetrazolium arabinose (TA) indicator agar plates on days 0, 1, 3, and every 3 days thereafter for the first 90 days to track the ratio of the Ara^−^ and Ara^+^ lineages. Most of the populations in the 1000-fold dilution treatment had fixed one of the marker states by day 90, and we subsequently plated them only every 15 days. Similarly, we plated the populations in the 100-fold treatment every 15 days from day 120 onward.

### Fitness Measurements

We sampled clones from each population after 90 and 150 transfers (Table S11). The clones sampled after 150 transfers were taken from the numerically dominant marked lineage at that point. We used clones sampled after 90 transfers from the same marked lineage to avoid confusion while performing competition assays; in most cases, the same lineage was dominant at both time points. These clones competed against the ancestral strain with the opposite marker state, which allowed us to enumerate them by plating on TA agar. Our methods were the same as described elsewhere for the LTEE (3, 14, 16, 39, 48) except in three respects: (i) the competitions were performed in static test-tubes (the same as in our evolution experiment), rather than in shaken flasks; (ii) the volume of the inoculum and the dilution factor varied, as described above and indicated for the reported assays; and (iii) the competitions ran for multiple days, with daily dilutions, in order to estimate small fitness differences with greater precision. In particular, we ran competitions with 2-fold dilutions for 12 days (12 generations), with 8-fold dilutions for 6 days (18 generations), and with 100-fold and 1000-fold dilutions for 2 days (∼13.3 and ∼20 generations, respectively). We then calculated relative fitness as the ratio of the realized (i.e., net) growth rate of the evolved competitor to that of its ancestral counterpart during the competition: *W* = ln (*D^n^* × *E_f_* / *E_i_*) / ln (*D^n^* × *A_f_* / *A_i_*), where *D* is the dilution factor, *n* is the number of transfers (i.e., the duration of the assay minus 1), *E_i_* and *E_f_* are the initial and final densities of the evolved competitor, and *A_i_* and *A_f_* are the corresponding densities of the ancestor.

### Statistical Analyses

We performed all the statistical analyses using the referenced tests in R version 4.3.2.

## ACKNOWLEDGMENTS

We thank Chitrak Banerjee, Michael Desai, Ben Good, Thomas LaBar, Charles Ofria, and Mike Wiser for valuable discussions; Neerja Hajela and Jessica Baxter for assistance in the lab; and the reviewers for their constructive suggestions on an earlier version of our paper. We acknowledge funding from the Yamada Science Foundation (2016-5042 to MI), the Japan Society for the Promotion of Science (201860528 to MI), the MSU Professorial Assistants Program (to JPD), the National Science Foundation (DEB-1813069 and DEB-1951307 to REL), the US Department of Agriculture (MICL12143 to REL), and the BEACON Center for the Study of Evolution in Action (NSF Cooperative Agreement DBI-0939454).

