## Supplemental Information for "Effects of periodic bottlenecks on the dynamics of adaptive evolution in microbial populations"

**This PDF file includes:**

- Supplementary text
- Figures S1 to S10
- Tables S1 to S11
- SI References

### Supplementary Information Text

#### Methods

**Numerical simulations.** We wrote and implemented the STEPS (for Serially Transferred Evolving Populations Simulation) program to simulate the trajectories for relative fitness and neutral marker ratios in evolving asexual populations that undergo periodic serial transfers (https://github.com/zachmatson/STEPS/releases/tag/v1.0.3). Each run tracks a growing population of genotypes as they replicate and mutate; the population undergoes a transfer bottleneck after it reaches a final size, $N_f$, which reflects resource limitation. For each genotype, the program tracks its fitness, number of individuals, neutral marker state, and parent genotype, which are saved after every transfer. From these data, we can then compute the trajectories of the marker ratio and mean fitness as well as track mutation dynamics.

For the growth phase between transfers, we simulate a fixed number of timesteps based on the dilution factor, $D$. During each timestep, the growth rate of a lineage is determined by its fitness. The number of timesteps is such that the total population size approximately doubles in each timestep, except for the final one. The program calculates the duration of the timesteps, such that the length of each timestep (except the last one) is scaled inversely to the population mean fitness. This scaling allows those lineages with above-average fitness to more than double during that timestep, while lineages with below-average fitness increase less than two-fold. The last timestep in each transfer cycle is implemented differently, such that the total population grows only enough to reach the fixed final size (rather than double).

Beneficial mutations can occur during any timestep. The expected number of mutations equals the product of the beneficial mutation rate, $\mu_B$, and number of individuals produced in that timestep, with sampling from a Poisson distribution (1). The mutations are randomly distributed among lineages, with the expected numbers proportional to their relative abundances. Each mutation creates a new lineage with fitness $W_{new}$. Following the model of Wiser et al. (2), $W_{new} = W\,(1 + s)$, where $W$ is the fitness of the parent genotype relative to the ancestor, and $s$ is the effect size of the new mutation. The fitness effects are drawn from an exponential distribution, $\alpha e^{-\alpha s}$, that depends on the parent genotype's fitness, as follows: $\alpha_{n+1} = \alpha_n\,(1 + g\,s_{n+1})$, where $s_0 = 1/\alpha_0$ is the mean of the initial distribution, $s_{n+1}$ is the effect size of the $n$+1 mutation, and $g$ governs the strength of diminishing-returns epistasis.

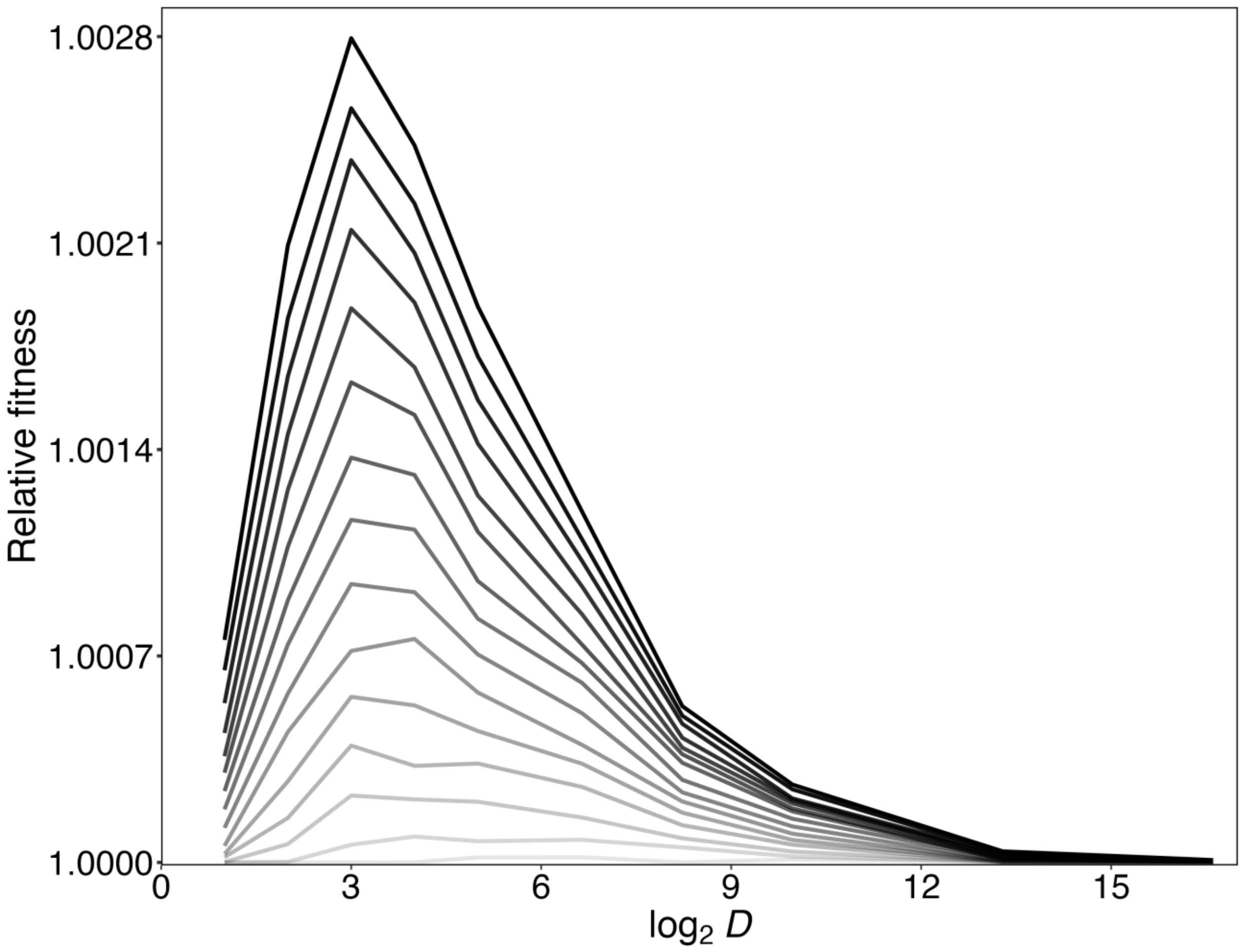


**Fig. S1.** Progression of mean relative fitness over time in the SSWM regime as a function of the dilution factor *D*. Data are from the same simulations shown in Figure 2 (main text), but they are presented here with time running from the bottom (100 transfers) to the top (1500 transfers). Over time, the profile stabilizes with the maximum fitness gains for $D = 8$ (i.e., $\log_2 D = 3$).

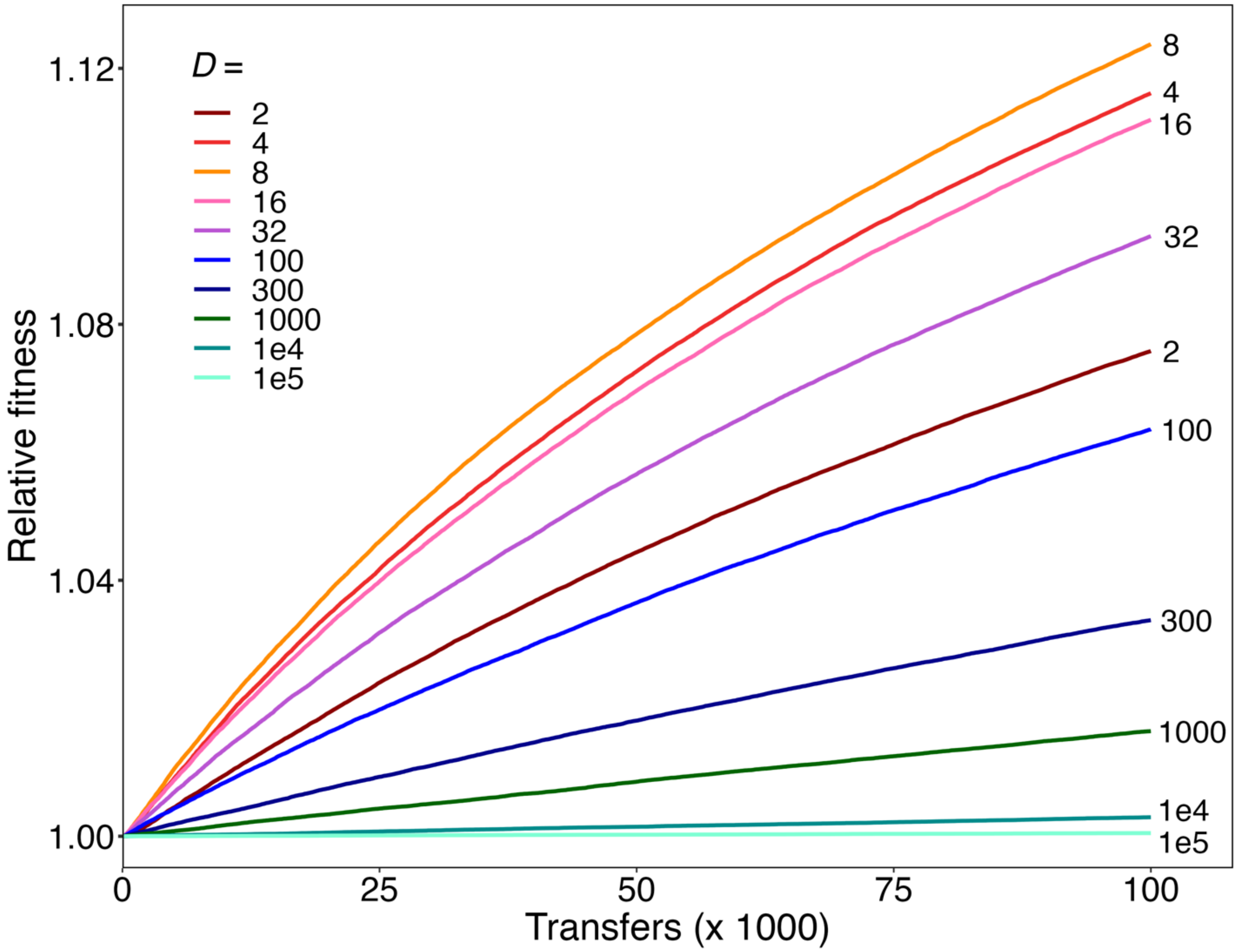


**Fig. S2.** Extended simulations showing trajectories of relative fitness for populations evolving under the SSWM regime with dilution factors ranging from 2-fold to $10^5$-fold. Each trajectory shows the grand mean of 10,000 runs for 100,000 transfers.

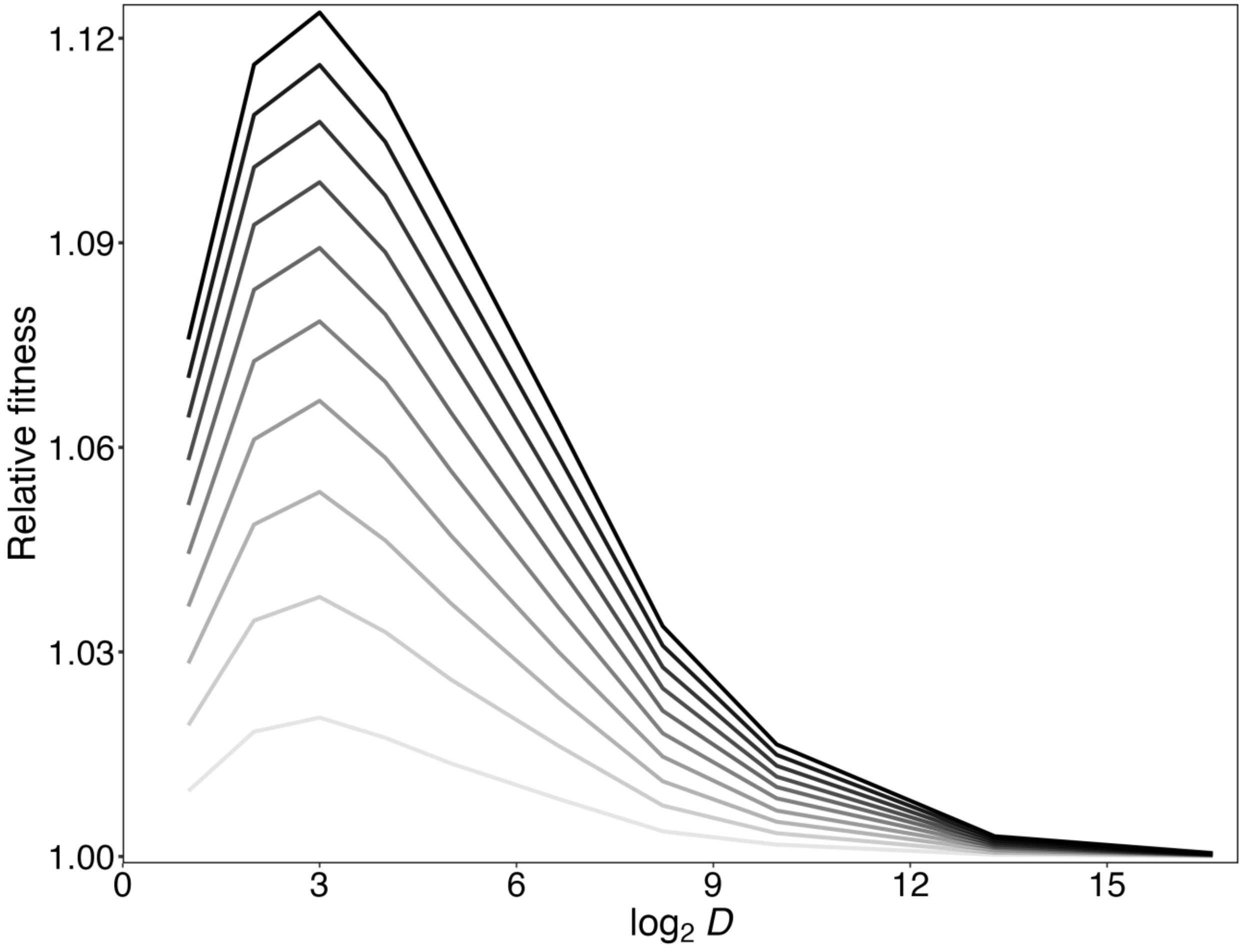


**Fig. S3.** Progression of mean relative fitness over time in the SSWM regime as a function of the dilution factor *D*. Data are from the same simulations shown in Figure S2, but they are presented here with time running from bottom (10,000 transfers) to top (100,000 transfers), thus extending the analysis shown in Fig. S1. The profile has clearly stabilized with the maximum fitness gains for $D = 8$ (i.e., $\log_2 D = 3$).

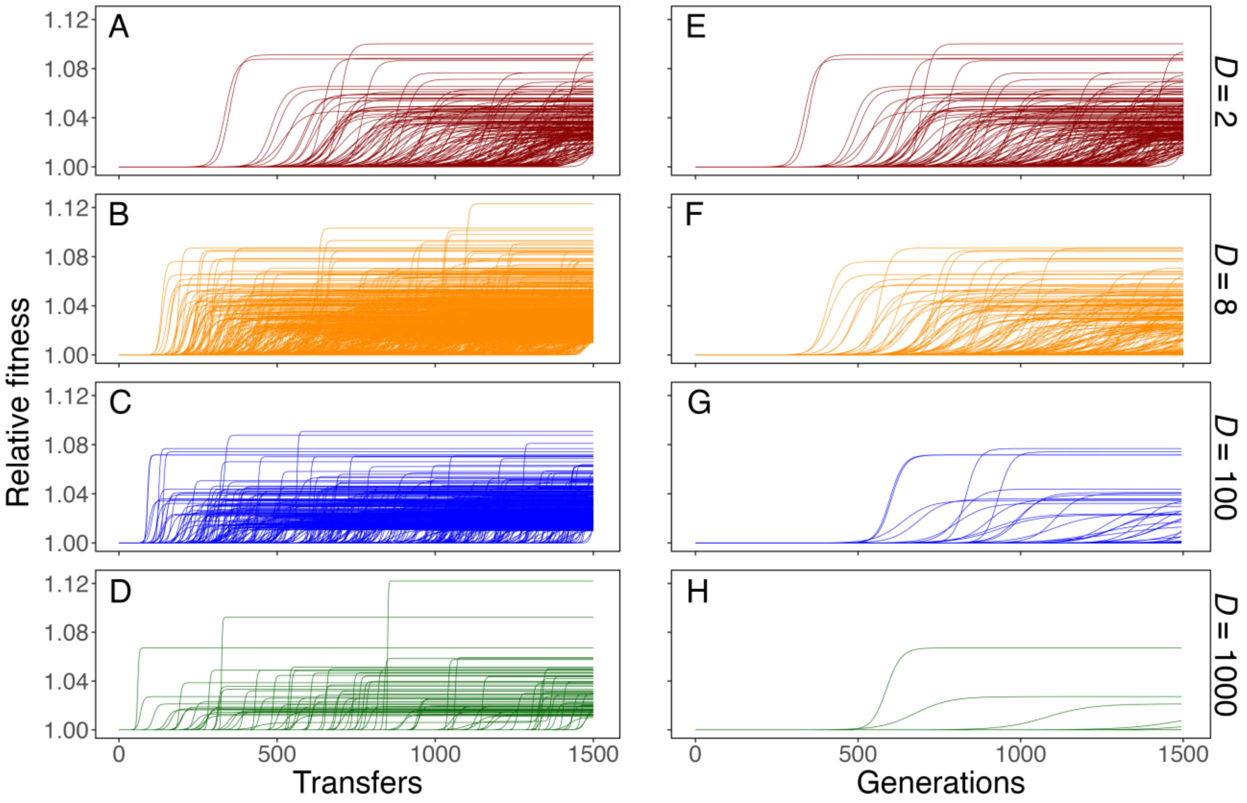


**Fig. S4.** Simulated relative-fitness trajectories with time shown for 1500 transfers (A-D) or 1500 generations (E-H) for 10,000 populations evolving under the SSWM regime with four dilution treatments. Most trajectories are flat lines (i.e., no fitness increase), because most populations in this regime had no beneficial mutations (Table S1). Note the steeper trajectories leading to similar fitness gains at the higher dilutions when plotted against transfers, but not when plotted against generations. Trajectories for the grand means are shown in Figure 2 (main text).

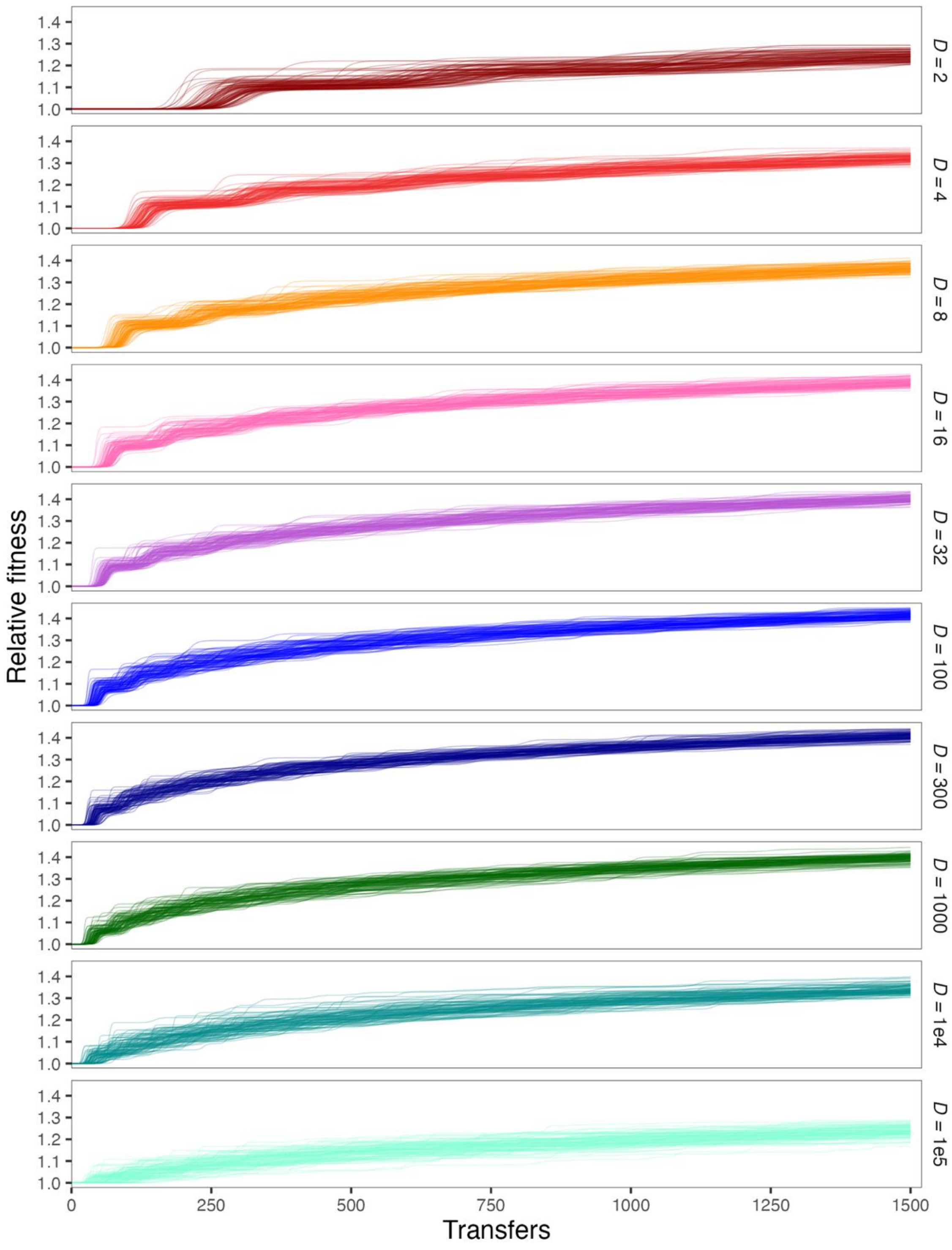


**Fig. S5.** Simulated fitness trajectories for 100 populations evolving under the SSSM regime with dilution factors ranging from 2-fold to $10^5$-fold. Figure 3 (main text) shows the grand means. Figs. S6 and S7 show high-resolution trajectories for the early transfers.

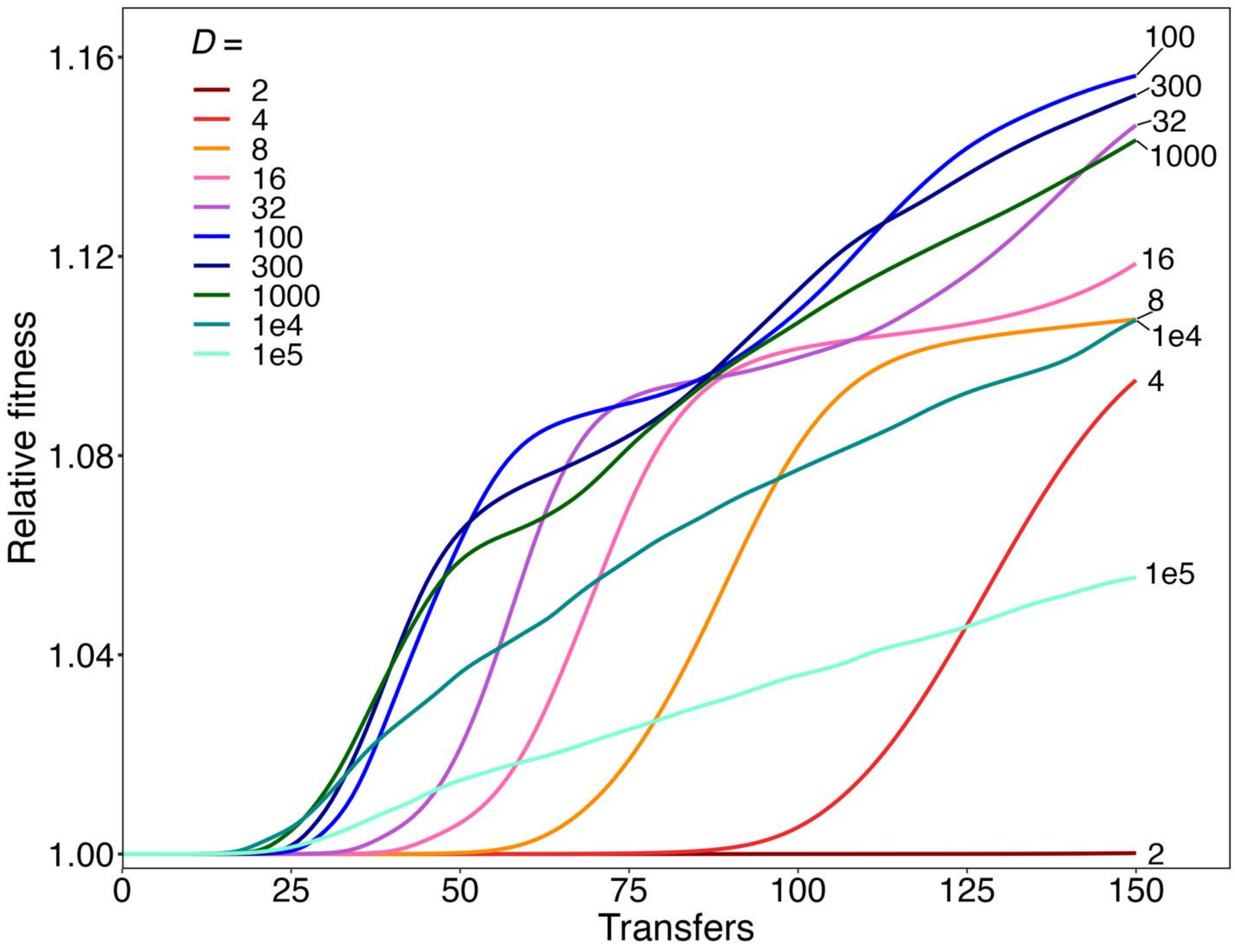


**Fig. S6.** High-resolution graph of the early phase of the simulated mean fitness trajectories for populations evolving under the SSSM regime with dilution factors from 2-fold to $10^5$-fold. Each trajectory shows the grand mean of 100 runs for the first 150 transfers only.

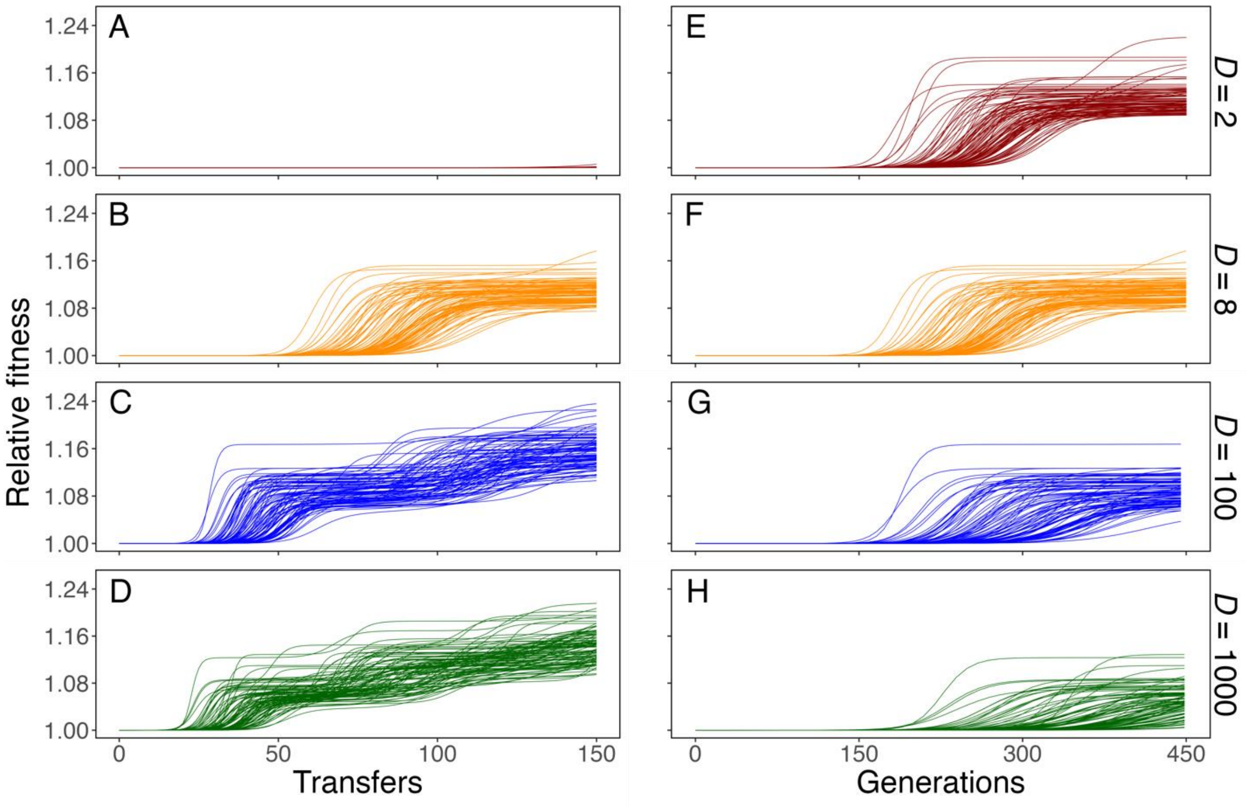


**Fig. S7.** Simulated fitness trajectories for 100 populations evolving under the SSSM regime with *D* = 2, 8, 100, and 1000 for 150 transfers (A-D) or 450 generations (E-H).

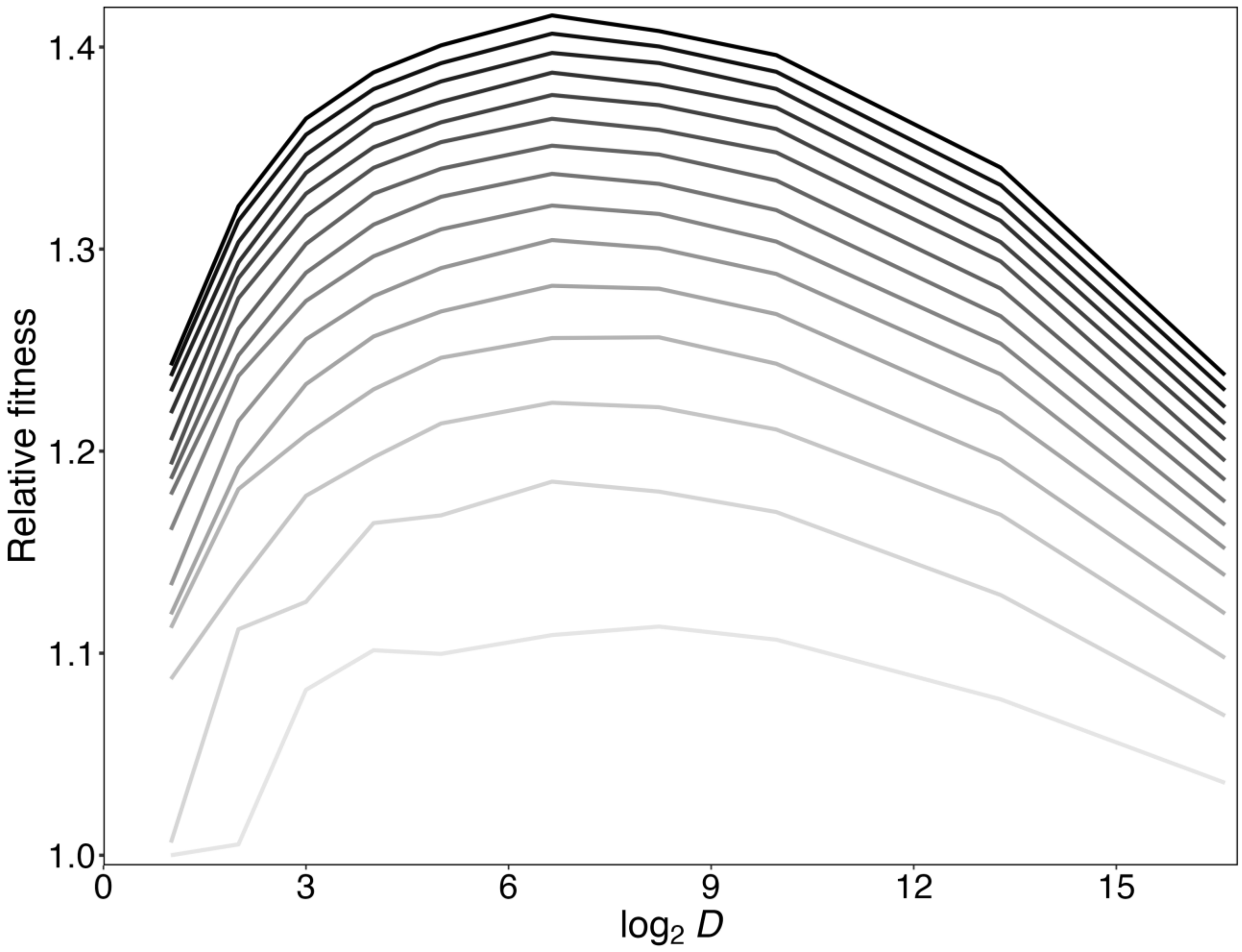


**Fig. S8.** Progression of mean relative fitness over time in the SSSM regime as a function of the dilution factor *D*. Data are from the same simulations shown in Figure 3 (main text), but they are presented here with time running from bottom (100 transfers) to top (1500 transfers). By 200 transfers, the profile has stabilized with the maximum fitness gains for $D = 100$ (i.e., $\log_2 D = 6.64$).

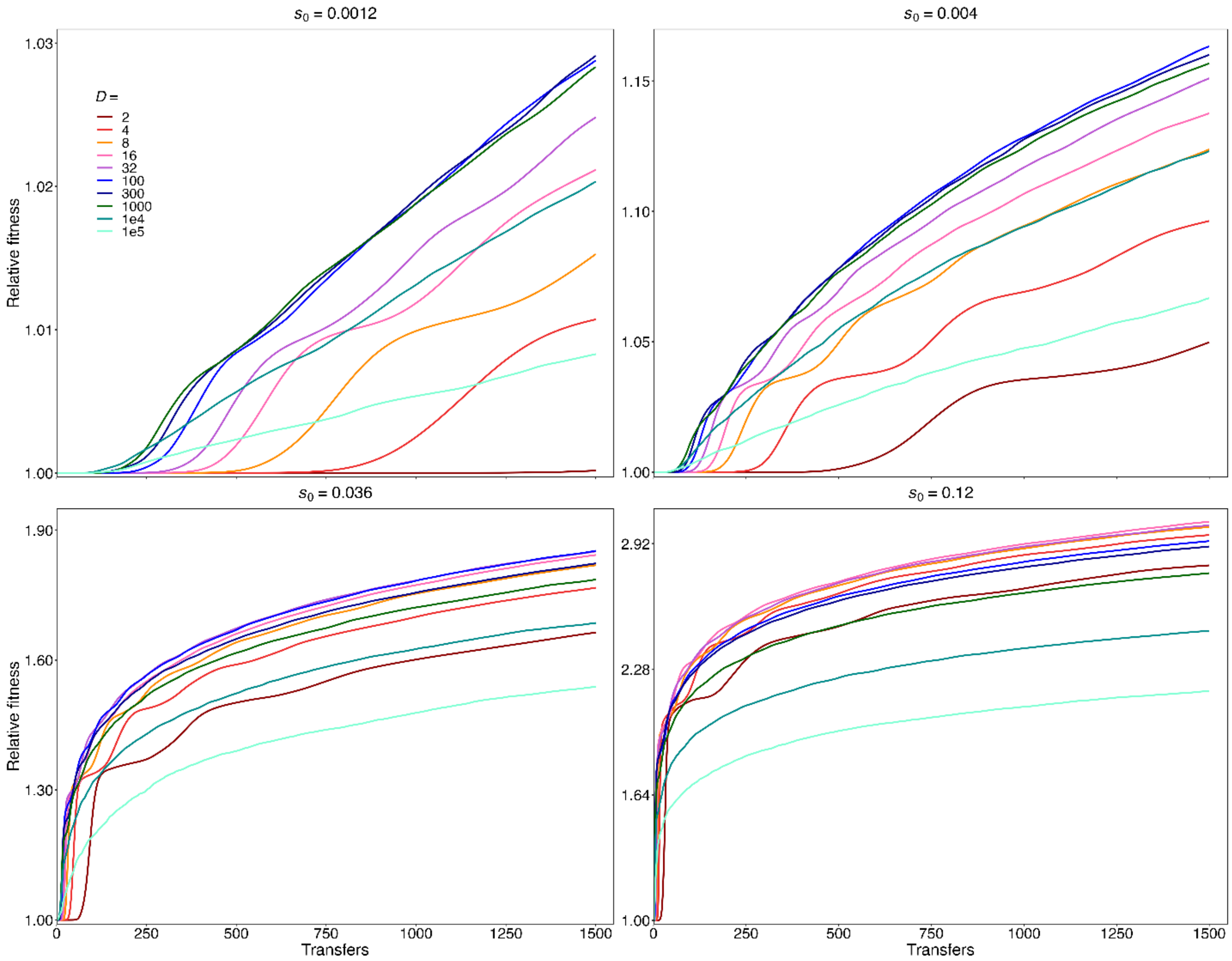


**Fig. S9.** Effect of changing the effect size of beneficial mutations on the dilution factor that maximizes fitness gains. The parameter $s_0$ represents the average effect size of a beneficial mutation in the ancestral genetic background. Figure 3 (main text) shows the grand mean fitness trajectory from 100 simulations for each of 10 dilution factors when $s_0$ = 0.012, which is the estimate based on the LTEE. The upper left and right panels show comparable trajectories when $s_0$ is reduced by 10-fold and 3-fold, respectively. The lower left and right panels show comparable trajectories when $s_0$ is increased by 3-fold and 10-fold, respectively.

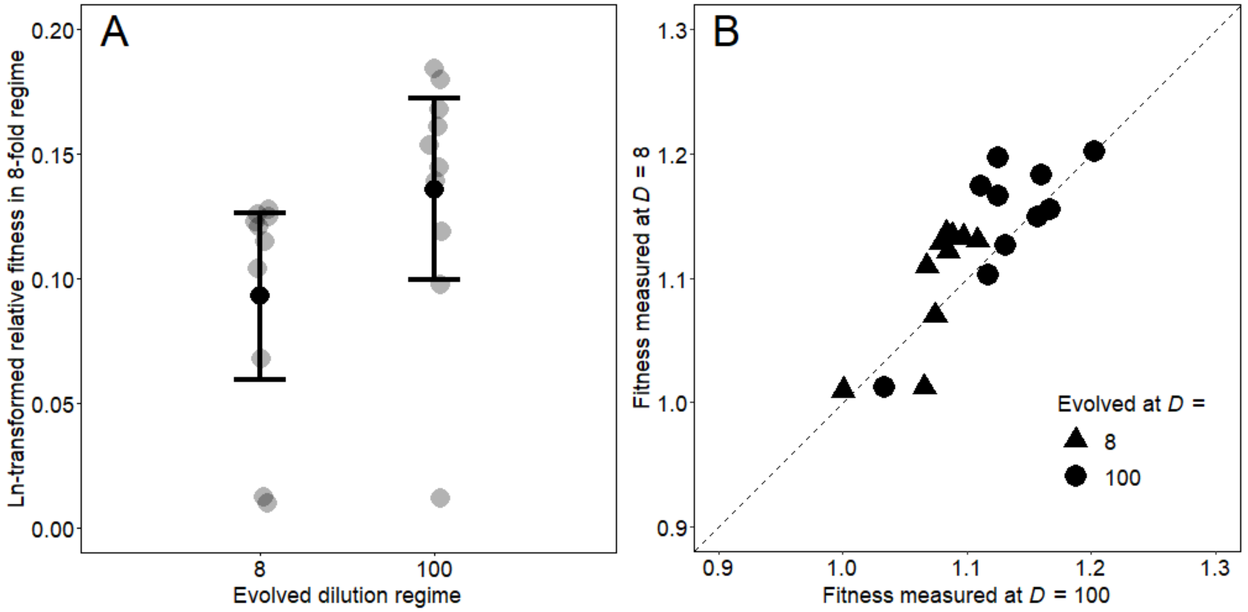


**Fig. S10.** Bacteria that evolved in the 100-fold dilution treatment were more fit than those that evolved in the 8-fold treatment. (A) Comparison of ln-transformed fitness values for bacteria that evolved in these two treatments, when both sets competed against marked ancestors with 8-fold dilutions. Black and gray symbols show means and replicate assays, respectively; error bars are 95% confidence intervals. The bacteria that evolved with 100-fold dilutions were more fit than those that evolved with 8-fold dilutions, even in the 8-fold treatment ($p$ = 0.0195, two-tailed Wilcoxon's signed-ranks test). (B) Correlation of fitness values measured with 8-fold and 100-fold dilutions for bacteria that evolved in those two treatments. The overall correlation is strong and highly significant ($r = 0.8218$, $p < 0.0001$).

**Table S1. Number of populations with surviving beneficial mutations and mean fitness after 1500 transfers in 10,000 simulated populations evolving under the SSWM regime.**

| Dilution factor (*D*) | Runs with surviving beneficial mutations | Mean fitness | *p* |
|---|---|---|---|
| 2 | 1030 | 1.0007545 | |
| | | | <0.0001 |
| 4 | 1447 | 1.0020922 | |
| | | | <0.0001 |
| 8 | 1525 | 1.0027946 | |
| | | | 0.0082 |
| 16 | 1225 | 1.0024313 | |
| | | | <0.0001 |
| 32 | 968 | 1.0018846 | |
| | | | <0.0001 |
| 100 | 555 | 1.0011901 | |
| | | | <0.0001 |
| 300 | 239 | 1.0005298 | |
| | | | <0.0001 |
| 1000 | 112 | 1.0002641 | |
| | | | <0.0001 |
| $10^4$ | 18 | 1.0000377 | |
| | | | 0.0162 |
| $10^5$ | 3 | 1.0000081 | |

We simulated 10,000 populations for each dilution treatment. We scored runs as having surviving beneficial mutations if the population's mean fitness was greater than unity at the end of the run. We compared the grand mean fitness of each treatment with the adjacent treatments (above and below) using two-tailed Welch's *t*-tests. All comparisons between adjacent treatments were significant ($p < 0.05$) after a table-wide sequential Bonferroni correction. Overall, mean fitness was highest with 8-fold dilutions, followed closely by the 16- and 4-fold dilutions, and it declined monontonically at both lower and higher dilution factors.

**Table S2. Mean fitness in simulated populations evolved under the SSSM regime after 1500 transfers with ten dilution treatments.**

| Dilution factor (*D*) | Mean fitness | St. dev. | *p* |
|---|---|---|---|
| 2 | 1.2424 | 0.0212 | |
| | | | <0.0001 |
| 4 | 1.3211 | 0.0155 | |
| | | | <0.0001 |
| 8 | 1.3645 | 0.0171 | |
| | | | <0.0001 |
| 16 | 1.3874 | 0.0156 | |
| | | | <0.0001 |
| 32 | 1.4008 | 0.0151 | |
| | | | <0.0001 |
| 100 | 1.4156 | 0.0146 | |
| | | | 0.0010 |
| 300 | 1.4079 | 0.0153 | |
| | | | <0.0001 |
| 1000 | 1.3960 | 0.0165 | |
| | | | <0.0001 |
| $10^4$ | 1.3403 | 0.0195 | |
| | | | <0.0001 |
| $10^5$ | 1.2376 | 0.0233 | |

We simulated 100 populations for each dilution treatment. We compared each treatment with the adjacent treatments (above and below) using two-tailed Mann-Whitney tests. All tests were significant ($p < 0.05$) after a table-wide sequential Bonferroni correction.

**Table S3. Change in mean fitness between transfers 1000 and 1500 in simulated populations with ten dilution treatments under the SSSM regime.**

| Regime (*D*) | Mean fitness | St. dev. | *p* |
|---|---|---|---|
| 2 | 0.0489 | 0.0146 | |
| | | | 0.0089 |
| 4 | 0.0454 | 0.0130 | |
| | | | 0.0237 |
| 8 | 0.0482 | 0.0112 | |
| | | | 0.4977 |
| 16 | 0.0471 | 0.0106 | |
| | | | 0.5567 |
| 32 | 0.0479 | 0.0104 | |
| | | | 0.0355 |
| 100 | 0.0511 | 0.0113 | |
| | | | 0.2423 |
| 300 | 0.0489 | 0.0106 | |
| | | | 0.3375 |
| 1000 | 0.0481 | 0.0118 | |
| | | | 0.3640 |
| $10^4$ | 0.0464 | 0.0125 | |
| | | | 0.0355 |
| $10^5$ | 0.0424 | 0.0165 | |

We simulated 100 populations for each dilution treatment. We compared each treatment with the adjacent treatments (above and below) using two-tailed Mann-Whitney tests. None of the comparisons were significant after a table-wide sequential Bonferroni correction.

**Table S4. Relative fitness of clones used to found aberrant populations in the marker-divergence experiments.**

| Clone pair | Mean fitness | Mean ln fitness | df | $t_s$ | $p$ |
|---|---|---|---|---|---|
| –3 vs +3 | 0.983 | –0.017 | 21 | –4.066 | 0.0006 |
| –5 vs +5 | 1.041 | 0.040 | 21 | 13.961 | <0.0001 |

We competed each member of a founding clone pair against the reciprocally marked ancestor, and the competition assays were paired. We calculated the relative fitness of a founding pair as the ratio of each clone's fitness relative to the ancestor. We replicated the paired competitions 22 times for each clone pair. We ran two-tailed paired *t*-tests on the ln-transformed fitnesses. Each comparison is highly significant and in the direction consistent with the initial divergence in the marker ratios (Fig. 4).

**Table S5. Comparison of times to divergence between the four dilution treatments in the experiment with bacteria.**

| Treatment comparison | *p* |
|---|---|
| 2-fold vs 8-fold | 0.0079 |
| 2-fold vs 100-fold | 0.0050 |
| 2-fold vs 1000-fold | 0.0050 |
| 8-fold vs 100-fold | 0.0122 |
| 8-fold vs 1000-fold | 0.0051 |
| 100-fold vs 1000-fold | 0.0146 |

We performed two-tailed Wilcoxon's signed-ranks tests to compare the times to divergence in the four dilution treatments. As explained in the text, two aberrant trajectories were excluded in each treatment; these tests are based on the remaining 10 trajectories in each treatment. All of the comparisons are significant even after a table-wide sequential Bonferroni correction ($p < 0.05$).

**Table S6. Mean time to divergence in simulated populations using ten dilution factors in the SSSM regime.**

| Dilution factor (*D*) | Mean *TD* | St. dev. | *p* |
|---|---|---|---|
| 2 | 316.27 | 114.8 | |
| | | | <0.0001 |
| 4 | 140.03 | 45.41 | |
| | | | <0.0001 |
| 8 | 101.27 | 36.02 | |
| | | | <0.0001 |
| 16 | 74.72 | 21.23 | |
| | | | <0.0001 |
| 32 | 63.51 | 21.25 | |
| | | | <0.0001 |
| 100 | 50.46 | 18.66 | |
| | | | 0.0122 |
| 300 | 45.42 | 15.05 | |
| | | | 0.0814 |
| 1000 | 43.10 | 16.39 | |
| | | | 0.8325 |
| $10^4$ | 42.98 | 14.91 | |
| | | | <0.0001 |
| $10^5$ | 61.37 | 30.42 | |

We simulated 100 populations for each dilution factor. The mean time to divergence, *TD*, is shown as the number of transfers. We compared the mean *TD* for each treatment with the adjacent treatments (above and below) using two-tailed Mann-Whitney tests. All tests were significant ($p < 0.05$) after a table-wide sequential Bonferroni correction except for the comparisons between 300- and 1000-fold and between 1000- and $10^4$-fold.

**Table S7. ANOVA of fluctuations in experimental marker ratios prior to the onset of divergence associated with selective sweeps.**

| Source | SS | df | MS | $F_s$ | $p$ |
|---|---|---|---|---|---|
| Treatment | 0.1278 | 3 | 0.0426 | 1.9677 | 0.1191 |
| Error | 5.9744 | 276 | 0.0216 | | |
| Total | 6.1022 | 279 | | | |

We calculated the absolute value of the $\log_2$-transformed change in the ratio of the two marked strains for the first 7 consecutive sample pairs (3-day intervals through day 21) for 10 populations in the 4 dilution treatments (excluding the two aberrant populations in each treatment). By day 24, some populations exhibited systematic divergence in marker ratios caused by selective sweeps, which thus set the limit of the data used in this analysis. The grand mean and standard deviation of these 280 early fluctuations were 0.1986 and 0.1479, respectively. The ANOVA shows no significant effect of the dilution treatment on the magnitude of the fluctuations.

**Table S8. Comparison of fitness measured in evolved and common environments of bacteria propagated for 150 transfers under four dilution treatments.**

| Treatment | Mean ln fitness Evolved | Mean ln fitness Common | df | $t_s$ | $p$ |
|---|---|---|---|---|---|
| 2-fold | 0.2138 | 0.0601 | 9 | 6.2641 | <0.0001 |
| 8-fold | 0.1351 | 0.0841 | 9 | 5.2474 | 0.0003 |
| 100-fold | 0.1249 | 0.1420 | 9 | –1.9668 | 0.0808 |
| 1000-fold | 0.1047 | 0.1326 | 9 | –3.2851 | 0.9953 |

We isolated clones at the end of the 150-transfer experiment from populations in the four dilution treatments. Each evolved clone competed against the reciprocally marked ancestral strain using both the same dilution treatment in which it evolved and the common 100-fold dilution treatment. We ran paired *t*-tests to compare the mean ln-transformed fitness values in the two environments. The tests were one-tailed in all cases except the 100-fold treatment, given the expectation that fitness would be higher in the environment where a population had evolved than in the common environment; the environments are identical for the 100-fold treatment, and so that test was two-tailed. Note that the reported *p*-value is high for the 1000-fold treatment because the difference was opposite to the expectation.

**Table S9. Comparison of final fitness of bacteria that evolved for 150 transfers under the 8-, 100-, and 1000-fold dilution treatments.**

| Treatment comparison | df | $t_s$ | $p$ |
|---|---|---|---|
| 8-fold vs 100-fold | 9 | –3.3905 | 0.0037 |
| 8-fold vs 1000-fold | 9 | –2.2608 | 0.0417 |
| 100-fold vs 1000-fold | 9 | 0.2752 | 0.7867 |

We competed evolved clones against reciprocally marked ancestors in the common environment (i.e., 100-fold dilution treatment), with 5-fold replication. We calculated the mean ln-transformed fitness value for each population, and we ran two-tailed *t*-tests to compare those values for the treatments shown.

**Table S10. Comparison of fitness changes between transfers 90 and 150 in bacteria that evolved in the 8-, 100-, and 1000-fold dilution treatments.**

| Treatment comparison | *p* |
|---|---|
| 8-fold vs 100-fold | 0.3750 |
| 8-fold vs 1000-fold | 0.5566 |
| 100-fold vs 1000-fold | 0.9219 |

We ran two-tailed Wilcoxon signed-ranks tests to compare the changes in mean ln-transformed fitness values from 90 to 150 transfers for the treatments shown.

**Table S11. List of bacterial strains used in this study.**

| Strain | Treatment | Replicate | Transfer | Ara marker* |
|---|---|---|---|---|
| REL 606 | ancestor | | | – |
| REL 607 | ancestor | | | + |
| MI 762 | 2 | 1 | 90 | – |
| MI 693 | 2 | 1 | 150 | – |
| MI 761 | 8 | 1 | 90 | – |
| MI 692 | 8 | 1 | 150 | – |
| MI 759 | 100 | 1 | 90 | – |
| MI 739 | 100 | 1 | 150 | – |
| MI 758 | 1000 | 1 | 90 | + |
| MI 689 | 1000 | 1 | 150 | + |
| MI 766 | 2 | 2 | 90 | + |
| MI 697 | 2 | 2 | 150 | + |
| MI 765 | 8 | 2 | 90 | – |
| MI 696 | 8 | 2 | 150 | – |
| MI 764 | 100 | 2 | 90 | + |
| MI 695 | 100 | 2 | 150 | + |
| MI 763 | 1000 | 2 | 90 | + |
| MI 694 | 1000 | 2 | 150 | + |
| MI 770 | 2 | 3 | 90 | + |
| MI 701 | 2 | 3 | 150 | + |
| MI 769 | 8 | 3 | 90 | + |
| MI 700 | 8 | 3 | 150 | + |
| MI 768 | 100 | 3 | 90 | – |
| MI 699 | 100 | 3 | 150 | – |
| MI 767 | 1000 | 3 | 90 | – |
| MI 698 | 1000 | 3 | 150 | – |
| MI 3 | all** | 3 | 0 | – |
| MI 15 | all** | 3 | 0 | + |
| MI 775 | 2 | 4 | 90 | + |
| MI 705 | 2 | 4 | 150 | + |
| MI 774 | 8 | 4 | 90 | + |
| MI 704 | 8 | 4 | 150 | + |
| MI 773 | 100 | 4 | 90 | – |
| MI 703 | 100 | 4 | 150 | – |
| MI 771 | 1000 | 4 | 90 | + |
| MI 702 | 1000 | 4 | 150 | + |
| MI 779 | 2 | 5 | 90 | – |
| MI 709 | 2 | 5 | 150 | – |
| MI 778 | 8 | 5 | 90 | – |
| MI 708 | 8 | 5 | 150 | – |
| MI 777 | 100 | 5 | 90 | – |
| MI 707 | 100 | 5 | 150 | – |

| | | | | |
|---|---|---|---|---|
| MI 776 | 1000 | 5 | 90 | – |
| MI 706 | 1000 | 5 | 150 | – |
| MI 5 | all** | 5 | 0 | – |
| MI 17 | all** | 5 | 0 | + |
| MI 784 | 2 | 6 | 90 | + |
| MI 713 | 2 | 6 | 150 | + |
| MI 783 | 8 | 6 | 90 | – |
| MI 712 | 8 | 6 | 150 | – |
| MI 781 | 100 | 6 | 90 | + |
| MI 711 | 100 | 6 | 150 | + |
| MI 780 | 1000 | 6 | 90 | – |
| MI 710 | 1000 | 6 | 150 | – |
| MI 788 | 2 | 7 | 90 | + |
| MI 752 | 2 | 7 | 150 | + |
| MI 787 | 8 | 7 | 90 | + |
| MI 716 | 8 | 7 | 150 | + |
| MI 786 | 100 | 7 | 90 | + |
| MI 715 | 100 | 7 | 150 | + |
| MI 785 | 1000 | 7 | 90 | + |
| MI 714 | 1000 | 7 | 150 | + |
| MI 792 | 2 | 8 | 90 | + |
| MI 721 | 2 | 8 | 150 | + |
| MI 791 | 8 | 8 | 90 | – |
| MI 720 | 8 | 8 | 150 | – |
| MI 790 | 100 | 8 | 90 | – |
| MI 719 | 100 | 8 | 150 | – |
| MI 789 | 1000 | 8 | 90 | + |
| MI 718 | 1000 | 8 | 150 | + |
| MI 796 | 2 | 9 | 90 | – |
| MI 725 | 2 | 9 | 150 | – |
| MI 795 | 8 | 9 | 90 | – |
| MI 724 | 8 | 9 | 150 | – |
| MI 794 | 100 | 9 | 90 | + |
| MI 723 | 100 | 9 | 150 | + |
| MI 793 | 1000 | 9 | 90 | – |
| MI 722 | 1000 | 9 | 150 | – |
| MI 800 | 2 | 10 | 90 | – |
| MI 729 | 2 | 10 | 150 | – |
| MI 799 | 8 | 10 | 90 | + |
| MI 728 | 8 | 10 | 150 | + |
| MI 798 | 100 | 10 | 90 | – |
| MI 727 | 100 | 10 | 150 | – |
| MI 797 | 1000 | 10 | 90 | – |
| MI 726 | 1000 | 10 | 150 | – |
| MI 804 | 2 | 11 | 90 | – |

| | | | | |
|---|---|---|---|---|
| MI 733 | 2 | 11 | 150 | – |
| MI 803 | 8 | 11 | 90 | – |
| MI 732 | 8 | 11 | 150 | – |
| MI 802 | 100 | 11 | 90 | – |
| MI 731 | 100 | 11 | 150 | – |
| MI 801 | 1000 | 11 | 90 | – |
| MI 730 | 1000 | 11 | 150 | – |
| MI 809 | 2 | 12 | 90 | + |
| MI 737 | 2 | 12 | 150 | + |
| MI 808 | 8 | 12 | 90 | + |
| MI 736 | 8 | 12 | 150 | + |
| MI 807 | 100 | 12 | 90 | + |
| MI 735 | 100 | 12 | 150 | + |
| MI 810 | 1000 | 12 | 90 | + |
| MI 754 | 1000 | 12 | 150 | + |

* All Ara$^-$ and Ara+ strains derive from *E. coli* REL606 and REL607, respectively.

** Founding clones for populations that showed aberrant marker-ratio trajectories (Fig. 4) and unequal initial fitness values (Table S4).
